# Multi-Omics Meta-Analysis Provides Insights into Reversible Phosphorylation During Arabidopsis Skotomorphogenesis

**DOI:** 10.1101/2025.10.22.683993

**Authors:** Iván Federico Berco Gitman, Cecilia Eugenia María Grossi, Denise Soledad Arico, María Agustina Mazzella, Rita María Ulloa

## Abstract

We developed a pipeline to standardize and integrate publicly available transcriptomic, proteomic, and phosphoproteomic datasets from *Arabidopsis thaliana* etiolated seedlings. The pipeline is broadly adaptable, as its database schema and processing scripts can be applied to datasets from any organism or experimental condition. We applied it to etiolated seedlings to uncover protein kinases and protein phosphatases relevant to early skotomorphogenic development, a stage in which seedlings grow underground and rely on precisely regulated signaling networks to ensure survival and successful emergence. This multi-omics approach revealed a comprehensive landscape of protein expression at this developmental stage; however, we observed widespread discrepancies between transcript and protein abundance, pointing to post-transcriptional and/or post-translational regulation. This cross-study minimizes experimental bias and allows the detection of phosphorylation events that may have been overlooked in single-condition studies. A search of kinase specific motifs revealed that RxxS motifs were significantly enriched among phosphorylated peptides in protein phosphatases and microtubule-associated proteins, suggesting regulation by calcium dependent protein kinases (CPKs). CPK3 and CPK9 were identified as the most relevant isoforms in the proteome and phosphokinome, supporting their central role in skotomorphogenesis. Our results underscore the importance of phosphorylation-mediated signaling in coordinating early seedling development and provide a resource for dissecting regulatory networks and candidate genes involved in skotomorphogenesis.

## Introduction

An important advantage of integrating omics datasets generated by independent laboratories is the possibility of drawing robust biological conclusions under a double-blind framework, where data producers and data analysts remain unaware of each other’s hypotheses and expectations. Despite the growing availability of public databases and large-scale datasets, the main limitation in omics research now lies less in data production, which has become increasingly accessible and scalable, and more in the systematic, unbiased analysis and integration of these massive datasets to extract meaningful biological insights. *Arabidopsis thaliana* is a predominant model system for studying plant molecular, cellular, and evolutionary biology. Its widespread use has led to the development of multiple public databases, software tools, and online platforms freely available to the plant community. The multi-omics analysis pipeline described in this manuscript integrates RNAseq, proteomic, and phosphoproteomic datasets from Arabidopsis etiolated seedlings. However, the database schema and processing scripts are broadly applicable to any organism or experimental condition. Independent datasets can be loaded and queried separately, while source-specific adjustments may be required due to the heterogeneous formats typical of proteomic and phosphoproteomic data. In this study, we focused on the protein kinases (PKs) and protein phosphatases (PPs) involved in reversible protein phosphorylation providing a broader perspective on the molecular players involved in this developmental stage.

Reversible protein phosphorylation and calcium (Ca^2+^) play a key role in the response of plants to environmental and endogenous stimuli (Batistič and Kudla, 2012; Bredow et al., 2021; Li et al., 2022; Thor, 2019). This dynamic post-translational modification can modulate protein conformation, activation state, turnover, subcellular localization, and interactions. It involves PK activities, which transfer the gamma phosphate from ATP to specific serine (S), threonine (T), or tyrosine (Y) residues, and PP activities responsible for the removal of phosphate groups (Vetoshkina and Borisova-Mubarakshina, 2023). The Arabidopsis genome is predicted to encode roughly 1,000 PKs and 150 PPs (Kerk et al., 2002; Wang et al., 2003). Calcium-dependent protein kinases (CPKs) are the main Ca^2+^ sensor-transducers in plants; these proteins possess EF-hand calcium-binding motifs and a S/T PK domain, enabling them to combine sensing and responding activities. This multigenic family, first described in *Arabidopsis thaliana*, comprises 34 members divided into four (I–IV) subgroups (Cheng et al., 2002). Members of this family exhibit substantial similarity except for the N-terminal and C-terminal (NT and CT) variable domains. The NT is critical for subcellular localization, as it contains consensus sequences for myristoylation and palmitoylation (Cheng et al., 2002; Hrabak et al., 2003) and is important for substrate recognition (Ito et al., 2010).

Several CPKs contribute to adult plant developmental programs in shoots, roots, pollen tubes, and root hair growth. Moreover, CPKs are involved in metabolic processes and signal transduction pathways triggered by environmental stresses such as cold, drought, high salinity, wounding, or pathogens (reviewed in Delormel and Boudsocq, 2019). However, our knowledge about the role of CPKs during early stages of plant development remains limited. Skotomorphogenesis is an early developmental phase that demands energy consumption and is critical for the establishment of seedlings beneath the soil. After reaching the surface, light induces photomorphogenesis mainly through phytochrome and cryptochrome photoreceptors, enabling seedlings to become autotrophic (Josse and Halliday, 2008; Sajib et al., 2023a, 2023b, Fox 2012). In *Pharbitis nil* seedlings, protein levels and activity of PnCDPK56 were high in darkness and reduced upon light exposure (Jaworski et al., 2011). In Arabidopsis, Ca^2+^ signaling together with the CPK isoforms AtCPK6 and AtCPK12 regulate phytochrome B nuclear translocation during dark-to-light transition (Zhao et al., 2023). These results suggest dynamic CPK activity during a critical phase for seedling survival and environmental adaptation.

## Materials and Methods

### Computational Pipeline Developed

A program implemented in C# (.NET, open source) handles data ingestion into a relational SQLite database and is operated via a console interface. Data from TAIR (GFF files, chromosomal FASTA sequences, and GO annotations), as well as from ScanProSite and TMHMM-2.0 (Krogh et al., 2001), and NOn-regular secondary structure (NORSp,) were loaded into the database. For transcriptomic input, the pipeline converts raw RNAseq counts and RPKM values into TPM. Proteomic and phosphoproteomic matrices were mapped to their corresponding genes based on peptide sequences when available.

Quantitative normalization was performed in R using linear regression models, with the results written back into the SQLite database. Data visualization was also managed in R, including heatmaps, phosphorylation diagrams, and phylogenetic trees (from NWK files), as well as all figures included in this work. Extracting and plotting any subset of genes required only minor R handling. All source code, scripts, and datasets used in this study are available on GitHub (https://github.com/ifgitman/OmicIntegrator) to ensure reproducibility and facilitate reuse by the community.

### Datasets and thresholds applied in this study

We analyzed public RNA-seq, (Hartmann et al., 2016; Martín and Duque, 2021; Pfeiffer et al., 2014; Yu et al., 2019) and proteomic (Kruse et al., 2020; Reichel et al., 2016; Zander et al., 2020) data performed with Arabidopsis Col-0 or Ler seedlings grown under dark conditions for 2 to 6 days in half strength MS at 22°C (Table S1). To normalize RNAseq data across the four experiments, TPM values for each of the 27,533 genes were first averaged. Then, a linear regression in logarithmic scale was performed for each experiment (ln y) using the previously calculated average (ln x) as the reference. This regression-based transformation adjusted the original data, allowing for the computation of a Standardized TPM (S-TPM) for each gene in each experiment as well as a Standardized Average TPM (SA-TPM). A gene was considered transcribed if it showed S-TPM values > 5 in at least three RNA-seq datasets, and not transcribed if S-TPM values were < 1 in at least three experiments (Fig. S1). We applied a stringent cut-off to minimize background noise. Genes with S-TPM values between 1 and 5 fall into an undefined zone, where transcriptional activity is ambiguous. Within this range, it is not possible to confidently determine whether the gene is genuinely expressed or not. Genes with SA-TPM>50 were classified as abundantly transcribed.

Protein abundance was estimated by summing the abundance of all peptides corresponding to each protein. To standardize the proteomic data from the three datasets, a linear regression on a logarithmic scale was performed between the protein abundance values in each experiment (ln[y) and the corresponding SA-TPM values (ln[x) for genes with detectable transcript levels. Standardized protein abundance (S-prot) equal to 0 was assigned to proteins that were not detected. The fitted values were then averaged to compute a Standardized Average protein abundance (SA-prot) for each gene. We also analyzed phosphoproteomic datasets from Arabidopsis seedlings grown in darkness (Arico et al., 2021; Zander et al., 2020). Phosphopeptides whose abundance exceeded the threshold in at least two replicates within each dataset were mapped to their corresponding proteins. Additionally, we queried the dataset from Kruse et al. (2020), which reports post-translational protein modifications during this developmental stage. Protein presence was confirmed when it was detected in at least two proteomes with S-prot>0 and with unequivocal identification of exclusive peptides or phosphopeptides (Fig. S2) while protein absence was confirmed when S-prot was 0 across all three proteomes. As in transcript analysis, we applied a stringent cut-off here as well. Proteins that did not meet the defined criteria were classified as uncertain, likely reflecting background signal or low-confidence detection. Proteins with SA-prot>50 were classified as highly abundant. *Bona fide* phosphoproteins were required to fulfill the protein presence criteria.

AtPKs were filtered using Gene Ontology (GO) ‘protein kinase activity’ (GO:0004672) term, AtPPs were filtered using terms ‘protein phosphatase’, or GO:0004721 ‘phosphoprotein phosphatase activity’ and GO:1903293 ‘phosphatase complex’ and manually curated, and microtubule-associated proteins (AtMAPs) were identified as described in Arico et al. (2024). CPK targets reported by PhosPhAt 4.0 experimental data were analyzed. Phylogenetic analyses were conducted using MEGA-X (Kumar et al., 2018). To identify PK recognition motifs within the Arabidopsis phosphoproteomes of etiolated seedlings, we searched for known phosphorylation consensus sequences reported by Wang et al. (2013) focusing on LxRxxS and RxxS for CPKs and Ca^2+^/CaMPKs; [S/T]P and Px[S/T]P for GSK-3, CDKs, and MAPKs (proline-directed motifs); and Sx[D/E], SDx[D/E], SDxED, and SExE for CK2 activity (acidic motifs). We analyzed their distribution among phosphopeptides to infer kinase activities.

Expression data for selected AtPKs, AtPPs, and members of the AtCPK family across different tissues were obtained from the Arabidopsis eFP Browser (Developmental Map dataset). In addition, their expression in Arabidopsis seedlings exposed to white, blue, and red light was evaluated using RNA-seq datasets from Yu et al. (2019) and Hartmann et al. (2016).

## Results

### Multi-Omic Datasets Reveal Widespread Post-Transcriptional Regulation in Etiolated Seedlings

Using a standardized approach to normalize and integrate publicly available RNA-seq, proteomic, and phosphoproteomic datasets, we examined the relationship between transcript and protein levels in Arabidopsis etiolated seedlings. To ensure reliable cross-study comparisons, only proteins and transcripts meeting stringent detection criteria were retained, at the risk of excluding some true positives. Analysis of four selected RNA-seq datasets showed that 14,564 genes of the 27,533 protein coding genes (53%) were transcriptionally active, with 3,692 exhibiting high transcript levels (SA-TPM>50; dots to the right of the green line in Fig. 1A, Table 1). However, only one third of these transcribed genes (4,777 out of 14,564) had corresponding proteins (Fig. 1A, Table 1). Notably, 21% of the highly transcribed genes (777 out of 3,692) lacked detectable protein levels (blue dots in Fig. 1A) suggesting that many transcripts were either not translated or that the corresponding proteins were rapidly degraded. GO analysis of genes with SA-TPM>100 but no protein detection (SA-prot=0) revealed significant enrichment for terms related to photosynthesis and chloroplast thylakoid membranes (Table S2). Conversely, among the 5,043 proteins confidently detected across the three proteomic datasets, 46 corresponded to transcripts with S-TPM<1 in at least three experiments (magenta dots in Fig. 1A, Table 1). These “stored proteins” were enriched for GO terms associated with post-embryonic development, carbohydrate metabolism, and nutrient reservoir activity (Table S3). In agreement, eFP Browser data indicate that 20 of these genes are predominantly expressed in dry or imbibed seeds (Fig. S3).

**Figure 1.**
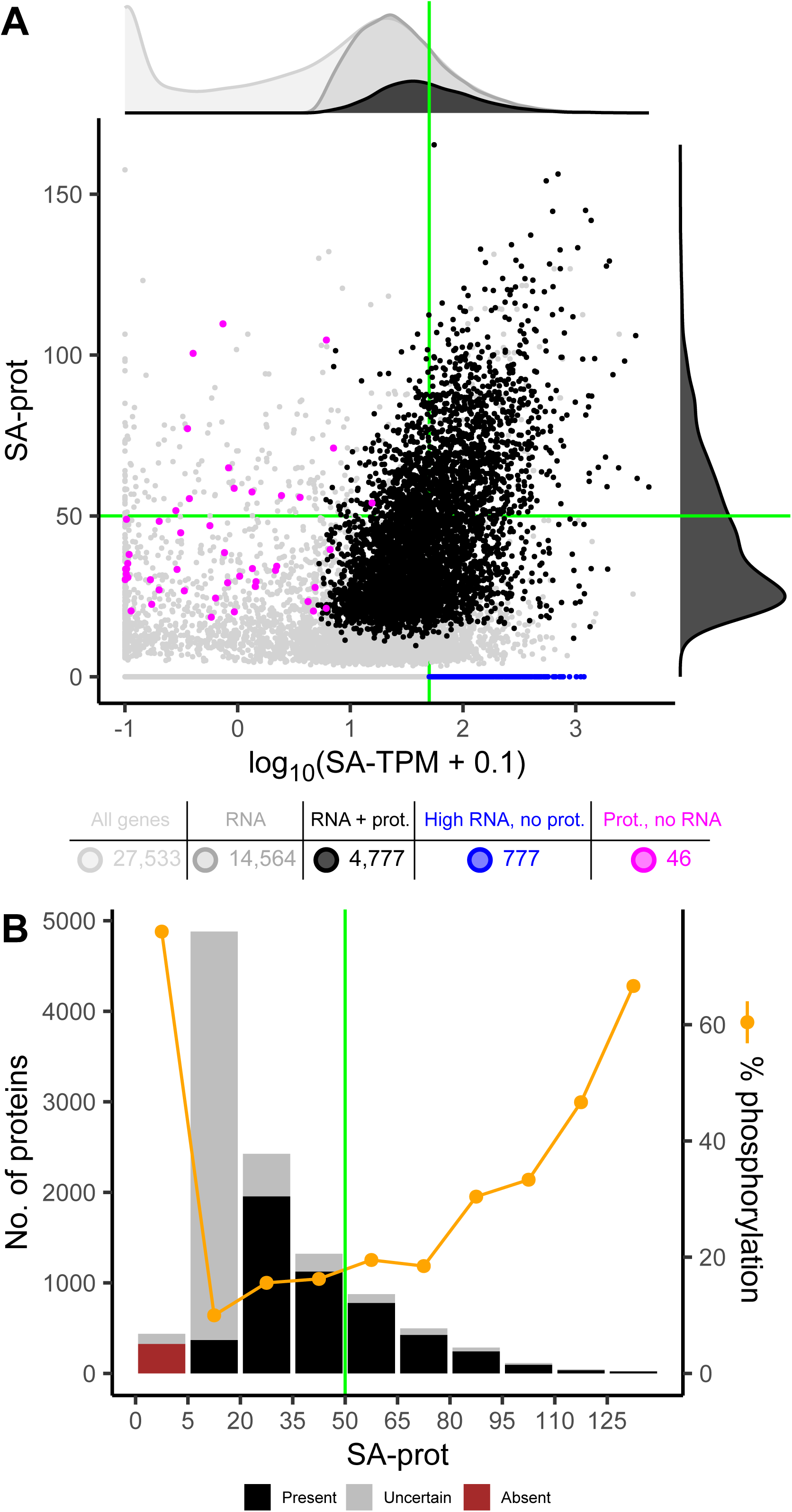
Multi-Omic Integration of Transcript, Protein, and Phosphoprotein Profiles in Dark-Grown Seedlings. **(A)** Scatter plot of transcript versus protein abundance for 27,533 protein-coding genes. Green lines mark the thresholds for high transcript or high protein abundance. 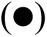 Proteins with corresponding transcript; 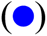 high transcript levels without corresponding protein; 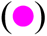 proteins without corresponding transcripts (stored proteins); 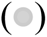 below thresholds or with ambiguous peptides. Gene counts per category are shown in the table below and their distributions are summarized in kernel density plots at the margins: top, all genes (light gray) transcribed genes (dark gray), and transcripts with corresponding protein (black); right: proteins with transcripts. **(B)** Protein distribution according to SA-prot values and percentage of phosphorylated proteins for each category. Black bars: highly confident proteins (detected in at least two datasets, with exclusive peptides), gray bars: uncertain proteins (detected in only one dataset and/or without exclusive peptides), brown bar: absent proteins (SA-prot=0). The green line marks the threshold for high protein abundance.

**Table 1.**
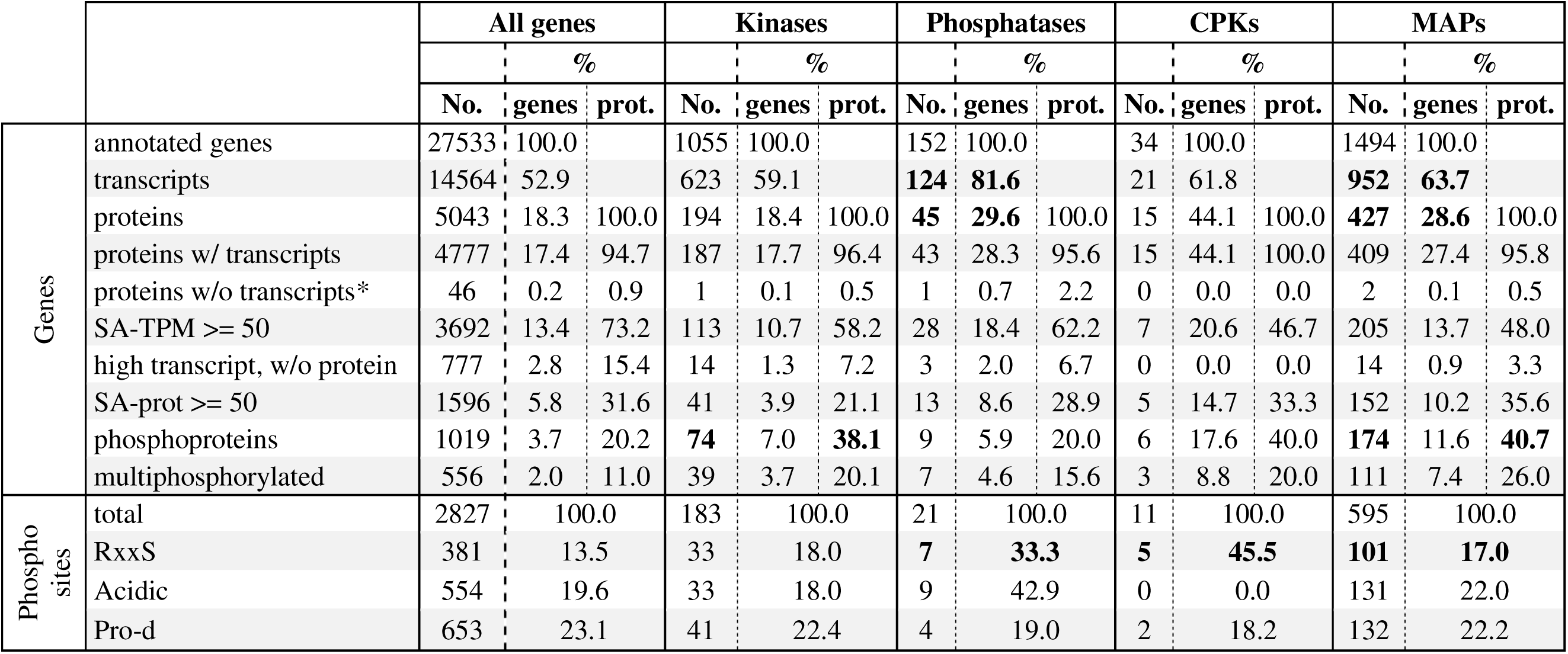
Summary of annotated genes, transcripts, and proteins in dark-grown *Arabidopsis thaliana* seedlings. The table presents the total number of annotated genes and their representation at the transcript and protein levels. Counts for PKs, PPs, CPKs, and MAPs are listed. Percentages (%) relative to the annotated genes or detected proteins in each gene set are shown. Phosphosites detected in the different protein sets were classified as RxxS (Ca^2+^/CaMPKs and CPKs), acidic (CK2) and Pro-d (GSK-3, CDKs and MAPKs). Categories that are significantly overrepresented (χ² test; p<0.05) compared to the whole genome, proteome, or phosphoproteome, are highlighted in bold. *proteins with uncertain transcript presence are not considered.

|  |  | All genes |  |  | Kinases |  |  | Phosphatases |  |  | CPKs |  |  | MAPs |  |  |
| --- | --- | --- | --- | --- | --- | --- | --- | --- | --- | --- | --- | --- | --- | --- | --- | --- |
|  |  | % |  |  | % |  |  | % |  |  | % |  |  | % |  |  |
|  |  | No. | genes | prot. | No. | genes | prot. | No. | genes | prot. | No. | genes | prot. | No. | genes | prot. |
| Genes | annotated genes | 27533 | 100.0 |  | 1055 | 100.0 |  | 152 | 100.0 |  | 34 | 100.0 |  | 1494 | 100.0 |  |
|  | transcripts | 14564 | 52.9 |  | 623 | 59.1 |  | **124** | **81.6** |  | 21 | 61.8 |  | **952** | **63.7** |  |
|  | proteins | 5043 | 18.3 | 100.0 | 194 | 18.4 | 100.0 | **45** | **29.6** | 100.0 | 15 | 44.1 | 100.0 | **427** | **28.6** | 100.0 |
|  | proteins w/ transcripts | 4777 | 17.4 | 94.7 | 187 | 17.7 | 96.4 | 43 | 28.3 | 95.6 | 15 | 44.1 | 100.0 | 409 | 27.4 | 95.8 |
|  | proteins w/o transcripts* | 46 | 0.2 | 0.9 | 1 | 0.1 | 0.5 | 1 | 0.7 | 2.2 | 0 | 0.0 | 0.0 | 2 | 0.1 | 0.5 |
|  | SA-TPM >= 50 | 3692 | 13.4 | 73.2 | 113 | 10.7 | 58.2 | 28 | 18.4 | 62.2 | 7 | 20.6 | 46.7 | 205 | 13.7 | 48.0 |
|  | high transcript, w/o protein | 777 | 2.8 | 15.4 | 14 | 1.3 | 7.2 | 3 | 2.0 | 6.7 | 0 | 0.0 | 0.0 | 14 | 0.9 | 3.3 |
|  | SA-prot >= 50 | 1596 | 5.8 | 31.6 | 41 | 3.9 | 21.1 | 13 | 8.6 | 28.9 | 5 | 14.7 | 33.3 | 152 | 10.2 | 35.6 |
|  | phosphoproteins | 1019 | 3.7 | 20.2 | **74** | 7.0 | **38.1** | 9 | 5.9 | 20.0 | 6 | 17.6 | 40.0 | **174** | 11.6 | **40.7** |
|  | multiphosphorylated | 556 | 2.0 | 11.0 | 39 | 3.7 | 20.1 | 7 | 4.6 | 15.6 | 3 | 8.8 | 20.0 | 111 | 7.4 | 26.0 |
| Phospho sites | total | 2827 | 100.0 |  | 183 | 100.0 |  | 21 | 100.0 |  | 11 | 100.0 |  | 595 | 100.0 |  |
|  | RxxS | 381 | 13.5 |  | 33 | 18.0 |  | **7** | **33.3** |  | **5** | **45.5** |  | **101** | **17.0** |  |
|  | Acidic | 554 | 19.6 |  | 33 | 18.0 |  | 9 | 42.9 |  | 0 | 0.0 |  | 131 | 22.0 |  |
|  | Pro-d | 653 | 23.1 |  | 41 | 22.4 |  | 4 | 19.0 |  | 2 | 18.2 |  | 132 | 22.2 |  |

Most reliably detected proteins (black bars in Fig. 1B) had SA-prot values between 20 and 50, while 1,596 were classified as highly abundant (SA-prot>50). By contrast, most uncertain proteins had SA-prot values between 5 and 20 (grey bars in Fig. 1B). We identified 1,839 phosphoproteins in at least one of the two available phosphoproteomic datasets from dark-grown seedlings (Arico et al., 2021; Zander et al., 2020), with 427 consistently detected in both. Strikingly, 325 phosphoproteins were not detected in any proteome (brown bar in Fig. 1B), including 37 with high transcript levels (SA-TPM>50), suggesting that phosphorylation could precede protein degradation (Table S4). Alternatively, detection of phosphopeptides without the corresponding proteins may reflect peptide enrichment, differences in detection sensitivity, or incomplete proteome coverage. After filtering according to protein presence, 1,019 phosphoproteins (with 2,827 phosphosites) were classified as bona fide (Table 1). At this developmental stage, phosphoproteins accounted for 20.2% of the proteome, a percentage that increased significantly (χ²=52.8, p<0.00001) among proteins with SA-prot>80 (Fig. 1B).

Large-scale analyses in plants have identified conserved kinase recognition motifs (Wang et al., 2013). Analysis of the phosphoproteome revealed that 23.1% of the phosphosites matched consensus motifs for GSK3, CDK, and/or MAPK activities; 13.5% for Ca²[/CaM-PKs and CPKs; and 19.6% for CK2 (Table 1). These assignments are based on shared consensus motifs that can be targeted by multiple kinases within a family or subfamily, and thus reflect combined rather than individual kinase activities.

By standardizing multi-omic datasets from etiolated seedlings we observed that many highly expressed genes lacked protein products, while others accumulated in the absence of corresponding transcripts. Together, these discrepancies between transcript and protein levels point to widespread post-transcriptional and post-translational regulation during skotomorphogenesis.

### Selective Expression of Protein Kinases in Etiolated Seedlings

The Arabidopsis genome encodes 1,055 genes annotated with protein kinase activity (GO:0004672). Of these, 623 were detected at the transcriptional level, with *MAPK17* showing the highest expression (SA-TPM=537, Supplementary Appendix). Only 187 (∼30%) had corresponding proteins identified in the proteomes of etiolated seedlings (Table 1), a transcript-to-protein ratio comparable to that observed for the entire transcriptome (∼34.6%). Among the *PK* genes not translated into detectable proteins, 14 showed high transcript abundance (SA-TPM>50, Table 1). Notably, this group includes six genes encoding calcineurin B-like (CBL)-interacting PKs: CIPK1/7/12/15, and SALT OVERLY SENSITIVE (SOS)3 interacting proteins SIP3/4, and a CPK-related protein (CRK2). Conversely, one PK gene (AT5G59680) encoding a Leucine-Rich Repeat Protein Kinase Family Protein (LRR-PK) was detected at the protein level despite having undetectable transcripts.

Of the 194 PKs identified in the proteome, 41 exhibited high protein abundance (Table 1), providing a snapshot of the PKs potentially active during skotomorphogenesis. Phylogenetic analysis (Fig. 2) revealed clustering into three major branches. One monophyletic group comprises 14 receptor-like kinases (RLKs), and three receptor-like cytoplasmic kinases (RLCKs) from the CARK family (Pti1-1/CARK7, Pti1-2/CARK8, Pti1-3/CARK5) which are known for their role in plant immunity (Forzani et al. 2011). Among the RLKs, six members belong to the LRR-III subfamily, while *FERONIA* (*FER*) and *HERCULES RECEPTOR KINASE 1* (*HERK1*) are classified within the CrRLK1L subfamily. Two Raf-like kinases, CONVERGENCE OF BLUE LIGHT AND CO2 (CBC2) and MIXED LINEAGE KINASE/RAF-RELATED KINASE 1 (MRK1), both members of the C7 subfamily, cluster within the same branch. Raf-like kinases are the closest relatives to RLKs in Arabidopsis (Shiu and Bleecker, 2001). CBC2, highly expressed in guard cells, mediates light-induced stomatal opening (Hiyama et al., 2017). Recent findings show that C7 Raf-like kinases associate with and are phosphorylated by CPK28, act redundantly in stomatal opening and immunity, and exhibit distinct substrate specificities from canonical MKKKs (Gonçalves Dias et al. 2024). Another branch includes four brassinosteroid signaling kinases (BSKs; RLCK XII) and two phytochromes (PHYA and PHYB), in separate subgroups. The third branch contains various soluble kinases: seven members of the SnRK/CPK superfamily (CPK3/4/5/9/21 and SnRK2.1/2.5), two of the AGC family (PHOT1 and PHOT2), CDC2 from the CMGC family, and PK3 which grouped distantly from three glycolysis-related phosphoglycerate kinases (PGKs).

**Figure 2.**
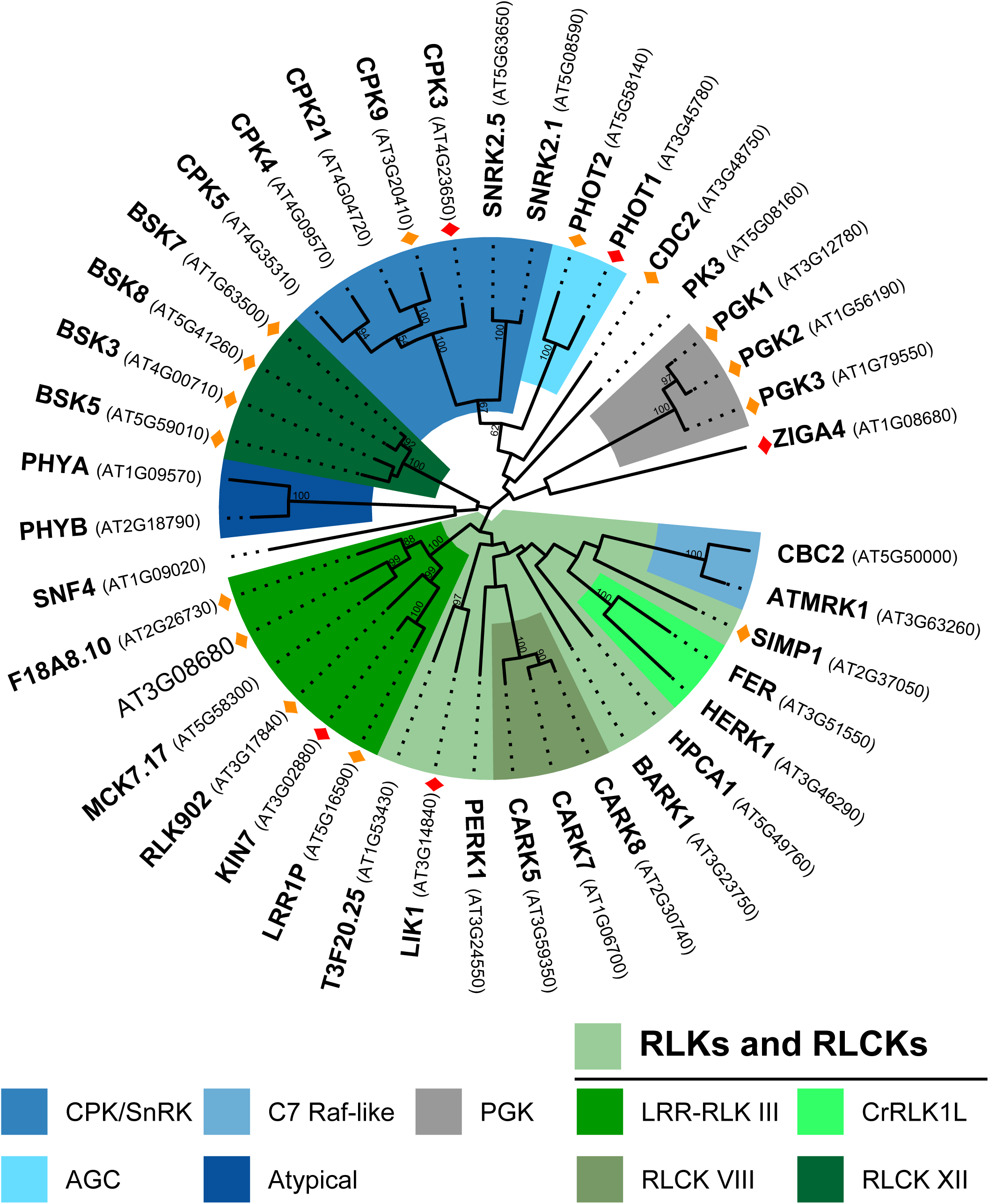
Phylogenetic analysis of 41 most abundant PKs (SA-prot>50) detected in etiolated seedlings. The inferred phylogeny was based on full-length protein alignments. Prominent PK families are color-coded. Bootstrap support values >50 (from 100 replicates) are shown at the nodes. Phosphorylated PKs are marked with 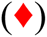 or 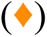 if detected in both or in one phosphoproteome respectively. SNF4 (a regulatory subunit of SnRK1), the zinc finger ARF GAP-like protein ZIGA4, and both phytochromes are not canonical kinases but were included due to their annotation with kinase-related GO terms.

Tissue expression analysis of these 41 PKs show that they are ubiquitously expressed except for mature pollen. Only CARK5 and 7 are strongly expressed in this particular tissue (Fig. S4). Transcriptomic analysis of seedlings exposed to different light treatments show that most PKs do not undergo significant transcriptional changes in response to light. Exceptions include the induction of RLK902, PHOT2, and PGK1, and the repression of SnRK2.5, BSK5, and PHYA (Fig. S4). Notably, SnRK2.5, which is highly expressed in dark-grown seedlings (SA-TPM: 156.96 and SA-prot: 61.51) shows markedly lower expression in other tissues, suggesting a specific role during skotomorphogenesis.

In conclusion, etiolated seedlings exhibit selective expression of PKs, and those involved in calcium signaling emerge among the most abundant soluble kinases.

### Phosphorylation Patterns of Protein Kinases in Etiolated Seedlings

Out of the 41 abundant kinases, 20 (48.8%) were phosphorylated (Fig. 2). Phosphorylated PKs were significantly overrepresented in etiolated seedlings (χ²=36.4, p<0.00001; Table 1), and the proportion increased further among highly abundant PKs, with 85.7% of those with SA-prot>80 being phosphorylated; far above the average phosphorylation rate for proteins of similar abundance (χ²=7.38, *p*=0.0066). In addition to these 20 phosphorylated PKs, 115 other PKs were identified in the phosphoproteome, including those with detectable and undetectable protein levels (Fig. S5). Phosphorylation events were particularly enriched in specific subfamilies, such as LRR-III kinases, BSKs, or MAP4Ks.

LRR-III kinases shared a conserved phosphosite in subdomain I, and three members were also phosphorylated in the CT (Fig. 3A). CT phosphorylation can create docking sites for downstream substrates in RLK signaling pathways (Pawson, 2004) and modulate substrate phosphorylation, as shown for BRI1 (Wang et al. 2005). In line with Mitra et al. (2015), CT phosphosites in these LRR-III RLKs may facilitate substrate recognition. The seven BSKs identified displayed conserved phospho-serine residues (pS, Fig. S6A) that are known targets of BRI1 and essential for activation (Tang et al. 2008), indicating that they are active in etiolated seedlings. Phosphorylated versions of the nucleocytoplasmic phosphatases BSL1 and BSL2 (Fig. S6A), proposed BSK targets (Belkhadir and Jaillais, 2015), were also detected. BSK5, linked to plant immunity (Majhi et al. 2019, Majhi et al. 2021), carried a pS329 site (Fig. S6A). The five MAP4Ks were phosphorylated at multiple NT or CT sites, several in the RxxS context, and consistently detected in both phosphoproteomes (TOT3, TOI4, and TOI5, Fig 3B). These PKs positively modulate hypocotyl growth under warm temperature in etiolated seedlings (Vu et al. 2021).

**Figure 3.**
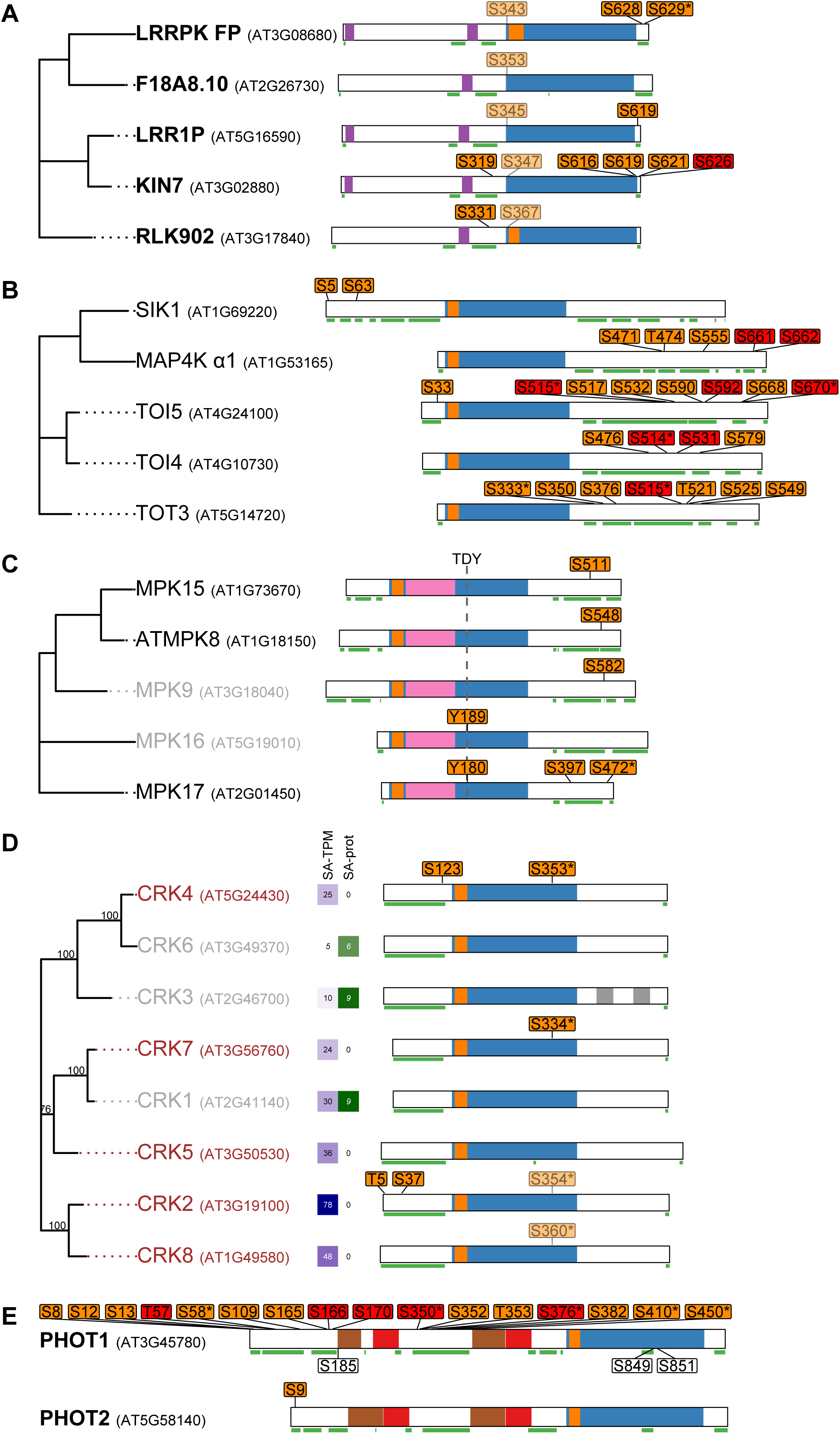
Phosphorylation patterns of LRR-III kinases **(A)**, MAP4Ks **(B)**, MAPKs **(C)**, CRKs **(D)** and Phototropins **(E)**. PK names in **bold**: high protein abundance, gray: uncertain presence, brown: not detected in any proteome. Domain architecture is based on ScanProSite and TMHMM predictions: 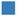KD, 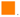ATP binding site, 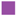transmembrane helix, 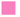MAPK domain, 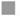EF hand, 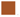PAS domain (Per-ARNT-Sim), 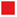PAC (PAS-associated C-terminal) motif. Disordered regions, based on PrDOS predictions, are <u>underlined in green</u>. pS/T: phosphosites detected in both phosphoproteomes, pS/T/Y: in only one dataset. Non-exclusive phosphopeptides are displayed with transparency. (*) Phosphorylations in an RxxS context. The ATP-binding site was absent in F18A8.10, LRR1, and KIN7. LRR1 and KIN7 belong to the atypical non-RD kinase subgroup and lack *in vitro* kinase activity (Chen et al., 2020). **(D)** SA-TPM and SA-prot values are indicated for CRKs. **(E)** In PHOT1, not phosphorylated residues S185, S849 and S851 are indicated.

The phosphokinome uncovered additional features. All identified MAPKs (8, 9, 15, 16, and 17) belong to group D1, characterized by a TDY activation motif, absence of a CD domain, and an extended CT region. CD domains contain acidic residues essential for interaction with a basic amino acid cluster in MAPKKs (Ichimura et al., 2002). MAPK16 and 17 exhibited a pY within the TDY motif, while MAPK8/9/15 and 17 were phosphorylated in the CT (Fig. 3C); these pS may provide an alternative MAPKKs docking mechanism. Among MAPKKs, only MKK2 (At4G29810) was confidently detected in the phosphokinome, while four others were present in the proteome at low abundance. Thus, MKK2 likely links the five MAPKs to the 21 MAPKKKs identified in etiolated seedlings (Fig. S5). These MAPKKKs belong to groups A, B and C; among them are three STY kinases involved in chloroplast differentiation (Lamberti et al., 2011). Five casein kinase 1-like (CKL) kinases were detected, all sharing conserved phosphosites in their CTs, a region responsible for determining substrate specificity (Fig S6 B).

Four CPK-Related Kinases (CRK2/4/7/8) were detected in the transcriptome and phosphokinome, but not in the proteome (SA-prot=0), suggesting that phosphorylation may negatively regulate protein stability during skotomorphogenesis (Fig. 3D). These CRKs shared phosphosites in a similar RxxS context ([A/G]<u>R</u>[T/S]E**p<u>S</u>**[A/G]IFR) within the catalytic domain. By contrast, CRK1/3/6 with SA-prot values between 6.2 and 9, were not phosphorylated.

The plasma membrane-associated PHOTs are blue light (BL) receptors that mediate phototropism, stomatal opening, chloroplast movement (Christie, 2007) and BL-induced increases in cytosolic free Ca²[ (Harada et al., 2003). PHOTs contain two NT LOV domains (PAC-PAS in Fig. 3E) separated by a hinge region and a C-terminal KD. Multiple phosphorylation sites have been identified within PHOTs, but only sites within the kinase activation loop are essential for all PHOT1-mediated responses (Inoue et al., 2008; Hart and Gardner, 2021; Christie et al., 2015). In etiolated seedlings, PHOT2’s unique phosphosite was restricted to its NT domain, while PHOT1 was phosphorylated at 16 residues (5 of which were detected in both phosphoproteomes) including nine in the NT and seven in the hinge region between the LOV domains (Fig. 3E). Residues S185, S849, and S851 were not phosphorylated under dark conditions, confirming their BL dependence, whereas pS58, pS170, pS350, pS376, and pS410, previously reported as BL-dependent (Inoue et al., 2008; Kinoshita et al., 2003; Salomon et al., 2003; Sullivan et al., 2008), were detected under dark conditions, as well as in other conditions reported in the PhosPhAt 4.0 database. Phosphorylation at S58, S350, S376, S410, and S450 occurred in the RxxS context suggesting modification by upstream PKs, as residues required for PHOT1 activity were not phosphorylated at this developmental stage (Fig. 3E). Hinge region phosphosites (pS350, pS376, pS410) are bound by 14-3-3 proteins (also known as General regulatory factors, GRFs) in etiolated seedlings (Kinoshita et al. 2003; Sullivan et al., 2009; Inoue et al., 2008). We identified eleven GRFs with high protein abundance (SA-prot 49.9–97), among which four were phosphorylated (Table S5).

In summary, we observed distinct phosphorylation patterns across PK families, including sites potentially linked to kinase activation (BSKs), substrate recognition (LRR-III RLKs and CKLs), protein–protein interactions (PHOT1–14-3-3, D1 MAPKs-MAPKKs), or protein degradation (CRKs). Phosphorylation frequently occurred in NT and CT domains, typically disordered regions, suggesting that these regions contribute to PK specificity and that phosphorylation may promote local structural ordering. Several phosphosites occurred in the RxxS motif, a known target of CPKs. Given their abundance at this developmental stage, we next examined the CPK family in detail.

### CPK3 and CPK9 are the main CPKs in dark-grown seedlings

Meta-analysis of the CPK family showed that 21 of the 34 AtCPK members were expressed above the transcript threshold, while only 15 were detected at the protein level in etiolated seedlings (Fig. 4A and B). The most abundant proteins included AtCPK3, AtCPK9, and AtCPK21 (subgroup II), AtCPK4, AtCPK5, and AtCPK6 (subgroup I), and AtCPK32 (subgroup III). Although AtCPK7 and AtCPK8 transcripts were highly abundant, their proteins could not be reliably detected (Fig. 4A). Most CPKs were ubiquitously expressed across tissues, excluding mature pollen, and were also detected in imbibed seeds, indicating their presence in the nascent seedling. AtCPK3, AtCPK5, and AtCPK32 maintained consistently high expression across tissues (Fig. S7). Transcriptomic datasets from dark grown seedlings exposed to white, blue and red light pulses (1 to 6 h) showed no major changes in CPK expression during dark-to-light transitions, suggesting that the same CPK set mediates calcium sensing during early development (Fig. S7).

**Figure 4.**
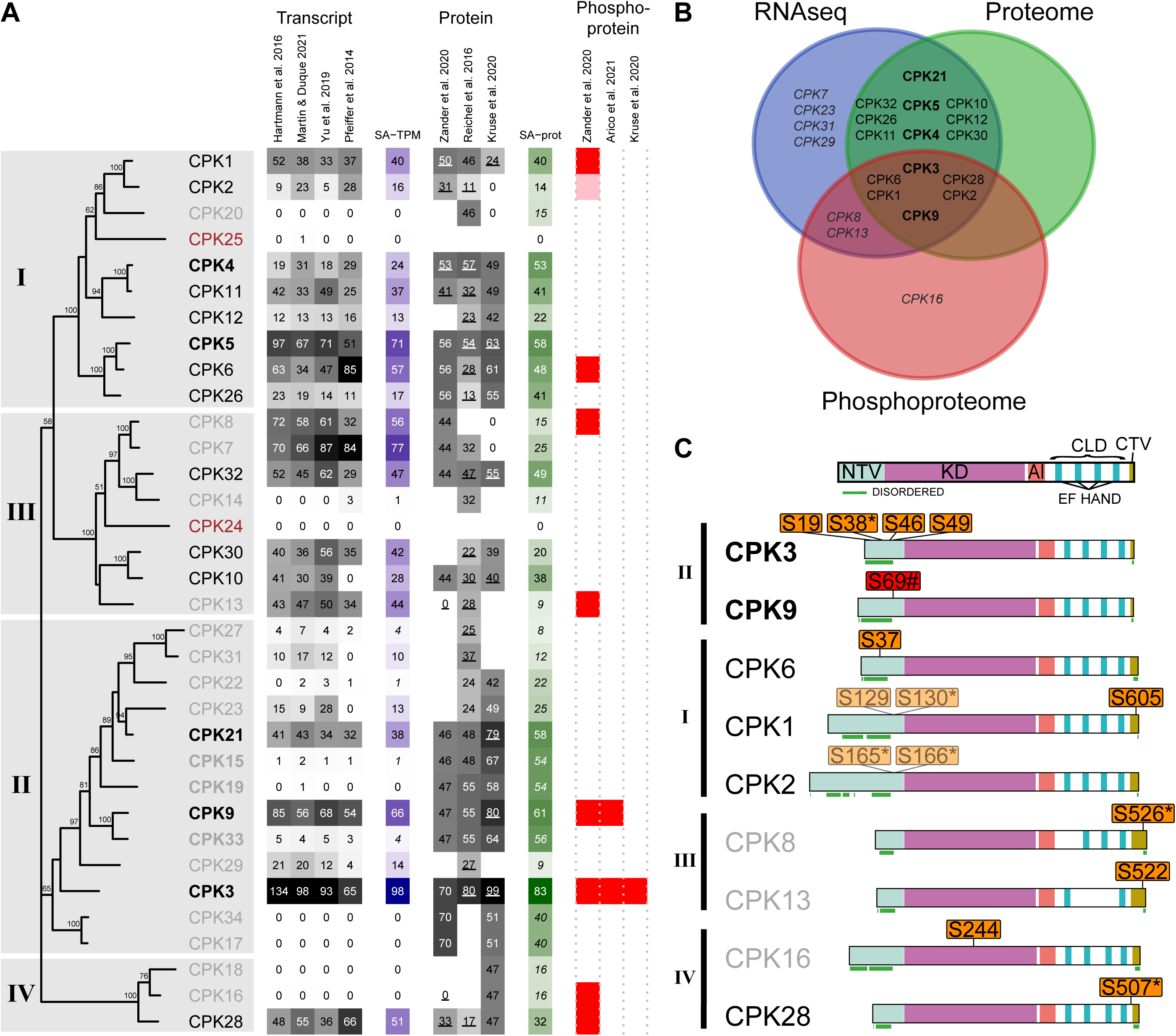
Meta-analysis of AtCPK transcripts, proteins, and phosphoproteins in dark-grown seedlings. **(A)** Phylogenetic tree of AtCPKs with four subgroups indicated. CPKs in **bold**: highly abundant, gray: uncertain presence; brown: absent. Transcript and protein abundances are shown as heat maps of normalized S-TPM and S-prot values (gray-scale, relative to the highest value), with SA-TPM or SA-Prot values on the right (blue or green scales, respectively). Numbers in *italics* in SA-TPM/SA-prot: uncertain or ambiguous; underlined in SA-prot: exclusive peptides. Presence in phosphoproteomes is indicated by red (exclusive phosphopeptides) or pink boxes (ambiguous phosphopeptides). Dataset authors are indicated in column headers. **(B)** Venn diagram of AtCPKs detected in transcriptome, proteome, and phosphoproteome datasets. **(C)** Domain structure of AtCPKs: NTV (N-terminal variable), KD (kinase domain), AI (autoinhibitory), CLD (calmodulin-like), CTV (C-terminal variable). Disordered regions are <u>underlined in green</u>. pS: detected in both phosphoproteomes, pS: in only one dataset. Non-exclusive phosphopeptides are displayed with transparency. (*) indicates phosphorylations in an RxxS context. (#) S69 detected by Zander et al. (2020, both replicates) and by Arico et al. (2021, one replicate).

CPKs are known to undergo autophosphorylation, which modulates their activity (Bender et al., 2017; Chaudhuri et al., 1999; Chehab et al., 2004; Ito et al., 2017; Oh et al., 2012; Saha and Singh, 1995; Sciorra et al., 2021; Ying et al. 2017). In etiolated seedlings, we identified phosphorylated forms of AtCPK1, AtCPK2, AtCPK3, AtCPK6, AtCPK8, AtCPK9, AtCPK13, AtCPK16, and AtCPK28 (Fig. 4 B and C). However, AtCPK8, 13 and 16 did not meet the criteria for *bona fide* phosphoproteins and *CPK16* transcripts were absent. Phosphorylated forms of AtCPK3 and AtCPK9 were detected in both phosphoproteomes and Kruse et al. (2020) also identified AtCPK3 as a phosphoprotein.

Most phosphosites were located within the NT domains of AtCPK1, AtCPK2, AtCPK3, AtCPK6, and AtCPK9 while additional phosphosites were identified in the CT domains of AtCPK1, AtCPK8, AtCPK13, and AtCPK28. NORSp analysis indicated that these NT and CT regions contain long stretches of disordered structure (NORS region, Fig. 4C). Several CPKs (AtCPK1/2/3/8/28) harbored pS in the RxxS context (Fig 4C). A χ² test comparing the frequency of phosphorylated RxxS motifs in the overall phosphoproteome (13.5%, 381 out of 2,827) with that in AtCPKs (45.5%, 5 out of 11) revealed a significant enrichment among AtCPKs (χ²=8.7, p<0.005) supporting the occurrence of intra- or intermolecular autophosphorylation events within this family.

The PhosPhAt 4.0 database reports 87 proteins as targets of one or more of the CPKs expressed in etiolated seedlings. Among these, we identified 47 in the proteome datasets and 18 in the phosphoproteomes. Notably, six proteins detected in both phosphoproteomes correspond to reported CPK3 or CPK9 targets, suggesting the functional activity of these kinases. Among them are the 14-3-3 proteins, GRF3/5. Furthermore, CINV1, RSZ22a, HA2, PP2C33, and AMT1;1 exhibit pS within the canonical RxxS motif (Fig. S8).

We also searched for phosphorylated proteins containing the RxxS motif, which is preferentially targeted by CPKs. RxxS motifs were significantly enriched among phosphorylated Microtubule Associated Proteins (MAPs, χ²=5, *p*<0.05; Table 1). Arico et al. (2024) showed that (MAPs) were overrepresented in the etiolated phosphoproteome. Reanalysis with the two available phosphoproteomes of dark-grown seedlings (Arico et al., 2021; Zander et al., 2020) confirmed this enrichment and revealed that 427 of the 1,494 MAPs encoded in the Arabidopsis genome are present at this developmental stage (χ²=97.6, *p*<0.00001), with 174 of them phosphorylated (χ²=97.4, *p*<0.00001). The cytoskeleton is dynamically regulated during early seedling development, contributing to differential cell growth and morphology in darkness (Walia et al., 2024). CPKs may modulate MAP activities through targeted phosphorylation events, highlighting them as potential upstream regulators of cytoskeletal dynamics at this stage.

Our analysis indicates that AtCPK3, and AtCPK9 are core components of the dark-grown seedling phosphokinome, consistently detected at the transcript, protein, and phosphoprotein levels.

### Most PPs are expressed and display broad Functional Diversity in Dark-Grown Seedlings

Protein phosphatases balance kinase activity; of the 152 *PPs* present in the Arabidopsis genome, 124 were transcriptionally active and 45 were detected with high confidence in the proteome of etiolated seedlings (Table 1). Interestingly, while the frequency of PKs (18.4%) closely matched the genome-wide protein detection rate (18.3%), PPs (29.6%) were significantly overrepresented (p=0.00034; χ²=12.8). This enrichment was primarily attributable to a significantly higher transcription rate (81.6%) compared to genome-wide average (52.9% χ²=49.93, p<0.00001), whereas their translation efficiency (36.3%) did not significantly differ from that of the full transcriptome (34.6%). Only one PP, Release of DOrmancy 5/Delay Of Germination 18 (RDO5/DOG18) was present in the proteome despite having undetectable transcripts, while three had high transcript levels (SA-TPM>50) but no protein: a phosphatidylglycerophosphate phosphatase 1 (AtPGPP1, At2G35680) involved in phosphatidylglycerol biosynthesis and photosynthetic function (Lin et al. 2016), a thylakoid associated phosphatase (TAP38, At4g27800), and PP2C.D6 (AT3G51370) that inhibits the Na^+^/H^+^ antiporter activity of SALT OVERLY SENSITIVE (SOS1) under non-salt-stress conditions (Table 1, Supplementary Appendix).

Sixteen members of the monomeric PP2C (PPM) family, along with several phosphatases from different subclasses of the PPP family, were identified (Fig. 5). Catalytic (C) subunits of PP2A (PP2A-4 and PP2A), PP4 (PPX1), and PP6 (FYPP1) were detected; these closely related enzymes are collectively considered PP2A-like phosphatases (Moorhead et al., 2007). All three structural A subunits of PP2A (RCN1, PP2AA2, and PP2AA3) and three regulatory B subunits (Bα, Bβ, and B’γ/PUX5, Fig. 5) were also present, consistent with the formation of multiple PP2A holoenzymes with distinct substrate specificities and localizations (Farkas et al., 2007). PP2AA proteins likely act as scaffolds assembling both PP2A and PP6 holoenzymes. FYPP1 interacts with A subunits (Dai et al., 2012), and together with SAL2–4, the SAPS-like regulatory subunits (Stefansson and Brautigan, 2006), may form a PP6-type heterotrimer. The regulatory subunit PP4R3, required along with PPX1 for miRNA biogenesis (Wang et al., 2019), was present as well. In addition, four PP1 isoforms (TOPP1, TOPP2, TOPP3, and TOPP8), one PP5 (PAPP5, a phytochrome-specific PP5; Ryu et al., 2005), one PP7 (MAIL3), and two BSLs (BSL1 and BSL2) classified as “Others” by Kerk et al. (2008) were identified.

**Figure 5.**
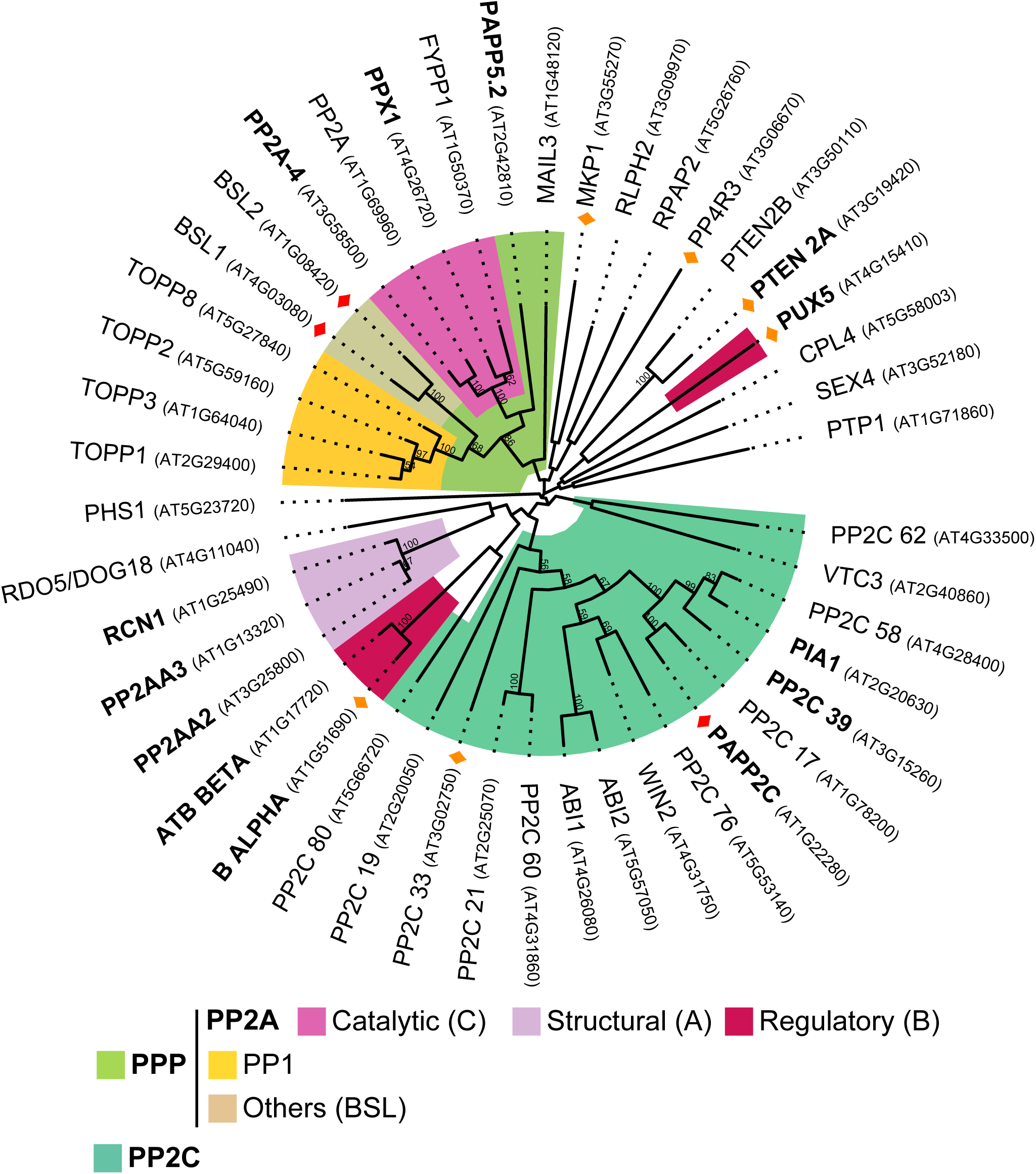
Phylogenetic analysis of 45 PPs detected in etiolated seedlings. The tree includes members of all major PP families. Sixteen members of the monomeric PPM family (PP2C) including VTC3, PP2C62, and 13 distributed among groups A (ABI1, ABI2), E (PP2C33), F1 (PIA1, PAPP2C, PP2C17/39/58), F2 (WIN2, PP2C76), I (PP2C21/60), K (PPC80), and L (PP2C19) (Bhaskara et al., 2019) were detected, together with 18 members of the PPP family including catalytic (C) subunits of PP2A (PP2A-4 and PP2A), PP4 (PPX1), and PP6 (FYPP1), regulatory and structural subunits of PP2A, 4 TOPPs (PP1), PAPP5 (PP5), MAIL3 (PP7), BSLs classified as “Others” (Kerk et al., 2008) and a PP4 regulatory subunit. Furthermore, the dataset included six members of the Class I PTP family (DSPs), two CTD phosphatases (CTD-PPs), and two protein tyrosine phosphatases (PTP and RLPH2). Bootstrap support values >50 (from 100 replicates) are shown at the nodes. Phosphorylated PPs are marked with 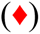 or 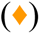 if detected in both or in one phosphoproteome respectively. PPs with high protein abundance (SA-prot>50) are indicated in **bold**.

PP2A subunits A, B, and C, along with PPX1, PAPP5, PTEN2A, and three PP2Cs (PIA1, PP2C39, and PAPP2C) exhibited high protein abundance suggesting that these enzymes play central roles in regulating protein phosphorylation turnover during skotomorphogenesis. Most of these PPs exhibited a constitutive expression pattern across all tissues, including mature pollen, with only six showing no detectable expression in it (Fig. S9).

Phosphorylation was detected in 20% of the PPs present in the proteome (Fig. 5 and S5). In addition to PAPP2C, a phytochrome-associated PP2C (Phee et al., 2008) and PP2C33, reported as potential CPK3 targets (Mehlmer et al., 2010), other PPs or PP regulatory subunits were found phosphorylated (Fig. 5). Of the 21 phosphosites identified across these nine PPs, most mapped to intrinsically disordered regions and six, including those in BSLs (Fig. S6A), in Bα and B’γ regulatory subunits of PP2A, in PP2C33 and in MKP1 (Fig. S10), matched the RxxS consensus motif. Phosphorylation at this motif was significantly overrepresented (p<0.01, χ²=7) compared with the global phosphoproteome of etiolated seedlings (Table 1).

In summary, PPs show higher transcript and protein expression than the genome-wide average, with members of multiple families (PP2Cs, DSPs, PPMs) represented, highlighting their broad regulatory potential in etiolated seedlings. Phosphorylation of PPs suggests they are themselves regulated post-translationally.

## Discussion

By integrating transcriptomic, proteomic, and phosphoproteomic datasets from etiolated Arabidopsis seedlings, we provide a comprehensive overview of gene expression, protein accumulation, and phosphorylation during early skotomorphogenic development. Despite differences in experimental parameters among datasets (Table S1), darkness is the dominant factor shaping the biological response. Our standardized approach enabled direct comparisons across omic layers, revealing marked discrepancies between transcript and protein levels. For example, several photosynthesis-related genes showed high transcript abundance but lacked detectable proteins, suggesting post-transcriptional and/or posttranslational regulation to prevent premature accumulation of photosynthetic machinery. Conversely, the persistence of proteins without detectable transcripts highlights the stability of proteins synthesized at earlier stages (Fig. S3).

As seedlings depend on limited energy reserves, efficient allocation between root and shoot growth is essential. The abundance of PKs and PPs in etiolated seedlings provides insight into the signaling toolkit that underpins environmental sensing and developmental decision-making. Notably, FER and HERK1 emerged as abundant RLKs with known roles in cell wall integrity sensing and growth regulation (Richter et al., 2017). FER additionally modulates ABA signaling, as well as brassinosteroid (BR) and ethylene responsiveness (Deslauriers and Larsen, 2010; Tang and Guo, 2025). The presence of photoreceptors such as PHYA, PHYB, and phototropins indicates that seedlings are pre-equipped to perceive light cues. Importantly, most PKs and PPs showed limited transcriptional changes upon light exposure (Figs. S4, S9), supporting a model in which their activity is primarily regulated at the posttranslational level, enabling a rapid and coordinated transition from skotomorphogenesis to photomorphogenesis.

Placing individual signaling pathways within this multi-omic framework reveals their physiological context. Members of the LRR-III subfamily of RLKs, as well as BSKs and CPKs, were among the most abundant and frequently phosphorylated PKs, supporting their central role in signal integration during skotomorphogenesis (Fig. 2). For instance, LRR1 and KIN7 regulate germination and cotyledon greening in ABA signaling (Xie et al., 2024), while BR signaling, crucial for hypocotyl elongation, activates BRI1/BAK1–BSK–BSL cascades (Belkhadir and Jaillais, 2015; Kalbfuß et al., 2022; Masselli et al., 2014). In etiolated seedlings, phosphorylation of BSKs at essential activation sites, together with BSL1/2 phosphorylation (Fig. S6A), indicates active BR signaling. While BSKs likely mediate BSL phosphorylation, CLE19–PXL1 signaling can phosphorylate BSLs in pollen (Wang et al., 2025) suggesting that multiple kinases may be implicated.

Within this signaling landscape, CPK3 and CPK9 stood out as core nodes of the etiolated phosphokinome, consistently detected across omic layers and associated with known targets (Fig. 4, Fig. S7). CPK6, also present in the three layers but with slightly lower abundance, has been previously linked to phyB signaling (Zhao et al., 2023), suggesting functional connections between light and dark signaling pathways. According to PhosPhAt 4.0, CPK3 and CPK9 contain 14 reported autophosphorylation sites, while CPK6 has seven, most within their NT domains. However, only a subset was phosphorylated in dark-grown seedlings (Fig. 4C), suggesting that the phosphorylation status of CPKs may vary depending on the plant’s developmental stage and/or environmental cues.

Under dark conditions, we observed phosphorylation of PHOT1 at RxxS motifs located in the hinge region, previously implicated in 14-3-3 (GRF) binding (Inoue et al., 2008; Kinoshita et al., 2003; Sullivan et al., 2009). CPKs and GRFs jointly regulate numerous aspects of plant biology; GRFs bind to RxxS phosphosites and modulate the activity, stability, subcellular localization, or complex assembly of their partners (Ormancey et al., 2017). CPKs can also phosphorylate GRFs, and we identified GRF3/5, reported as potential targets of CPK3 (Mehlmer et al., 2010), in both phosphoproteomes (Fig. S8). Moreover, a GRF has been reported to interact with AtCPK3 preventing its degradation (Lachaud et al., 2013, Ormancey et al., 2017). Given the RxxS phosphosites in the hinge region of PHOT1 and prior reports of GRF binding to PHOT1 in darkness (Kinoshita et al. 2003), we propose that CPK3 and these GRFs might regulate PHOT1 during skotomorphogenesis. In dark-grown seedlings, PHOT1 accumulates mainly in the hypocotyl hook and elongation zone (Sakamoto and Briggs, 2002; Wan et al., 2008) and associates with plasma membrane nanodomains together with AtRem (Xue et al., 2018). As remorin phosphorylation regulates nanodomain organization (Perraki et al., 2018), the multiple phosphosites detected in the disordered N-terminal region of PHOT1 (Fig. 3E) may underlie its targeting to specific membrane regions in the dark.

Phosphorylation of PP2A regulatory subunits at RxxS motifs (Fig. S10) suggests CPK-mediated control of phosphatase activity. Since PP2A specificity depends on its B subunits, such modifications could modulate complex assembly or function. Similar regulation has been described in rice (Ábrahám et al., 2015; Máthé et al., 2023), and together with the known CPK3 targets PAPP2C and PP2C33 (Mehlmer et al., 2010), these findings point to a dynamic interplay between CPKs and phosphatases in darkness. Additionally, the overrepresentation of phosphorylated MAPs (Arico et al., 2024) underscores the dynamic regulation of the cytoskeleton during early development. Many of these MAPs were phosphorylated at RxxS sites, further implicating CPKs as candidate regulators. Notably, AtCPK3 has been shown to control actin cytoskeleton organization during pathogen responses (Lu et al., 2020), raising the possibility that similar mechanisms operate during skotomorphogenesis to influence cell growth and morphology.

## Conclusions

We developed a pipeline to standardize and integrate transcriptomic, proteomic, and phosphoproteomic datasets from *A. thaliana* etiolated seedlings, which is broadly adaptable to other organisms or experimental conditions. During skotomorphogenesis, this integrative framework reveals dynamic changes in gene expression, protein abundance, and phosphorylation states that collectively shape the etiolated growth program. The results include findings that validate previous reports as well as novel or divergent observations, expanding current knowledge of signaling networks in dark-grown seedlings. Our analyses identify AtCPK3 and AtCPK9 as central regulators of reversible phosphorylation in darkness. Future studies should elucidate the specific contributions of these CPKs and their downstream phosphorylation targets to the regulation of skotomorphogenesis.

## Acknowledgements/funding details

RMU and MAM are members of CONICET. CEMG is a post-doctoral fellow of FONCYT, and IFBG is a doctoral fellow from CONICET. We acknowledge the support of CONICET and FONCYT through the following grants: PIP-CONICET 2018 to MAM and RMU, PICT 2021 0440 to RMU and PICT2018 0746 to MAM.

## Contributions

RMU and MAM conceived research plans, performed the meta-analysis and wrote the manuscript. IFBG developed the pipeline. IFBG, CEMG, and DSA conducted bioinformatic and statistical analysis. The authors declare no conflicts of interest.

## Data availability

Data is available at GitHub (https://github.com/ifgitman/OmicIntegrator).

## Supplementary material

**Table S1. Datasets analyzed in this work.** Experimental conditions reported by the original studies are shown; empty cells indicate unavailable data.

**Table S2. GO term analysis of 288 genes with very high transcript levels (SA-TPM>100) but no detectable protein (SA-prot=0) revealed a significant enrichment in photosynthesis-related genes**. Enrichment was assessed relative to the full transcriptome of etiolated seedlings (14,564 genes). MF: molecular function, BP: biological process, CC: cellular component.

**Table S3. GO term analysis of 46 proteins with undetectable transcript levels (S-TPM<1 in at least three experiments) revealed a significant enrichment in post-embryonic development, carbohydrate catabolism, and nutrient reservoir activity.** Enrichment was assessed relative to the whole genome of etiolated seedlings (27,533 genes). MF: molecular function, BP: biological process, CC: cellular component.

**Table S4.** List of 37 phosphoproteins with high transcript abundance (SA-TPM>50) but no detectable protein levels in any of the proteomic datasets. Phosphosites identified in both phosphoproteomic studies or only in one of them are indicated in separate columns.

**Table S5.** List of the 13 genes encoding for 14-3-3 proteins (GRFs).

**Figure S1. Normalization of RNA-seq data from four Arabidopsis dark-grown seedlings datasets.**

**(A)** Distribution of TPM values for 27,533 genes across four datasets. **(B)** Logarithmic linear regressions for each dataset, plotting the natural logarithm of the mean TPM value per gene across datasets (y-axis) against the natural logarithm of the TPM value for the same gene in the corresponding dataset (x-axis). Pearson correlation index is indicated. **(C)** Standardized TPM values (S-TPM) were obtained by applying a regression-based transformation to each gene in each dataset. The red line indicates the threshold used to consider transcript presence (S-TPM>5 in at least three experiments). The black line indicates the threshold used to infer absence of transcription (S-TPM values < 1 in at least three experiments). **(D)** Standardized average TPM (SA-TPM) values computed as the mean of the S-TPMs across all datasets. The green line indicates the threshold for highly abundant transcripts (SA-TPM>50).

**Figure S2. Normalization of proteomic data from three Arabidopsis dark-grown seedlings datasets.**

**(A)** Distribution of protein abundance across three datasets. **(B)** A linear fit on a logarithmic scale was performed between the protein abundance values in each experiment and their corresponding SA-TPM for genes with detectable transcript levels. **(C)** Distribution of the fitted values (S-prot) across the three datasets. **(D)** The fitted values were averaged to compute a standardized average protein abundance (SA-prot) for each gene. The blue line indicates the threshold for highly abundant proteins (SA-prot>50).

**Figure S3. Tissue expression pattern of the 46 proteins with undetectable transcript levels.** Genes in **bold**: highly abundant proteins (SA-prot values shown in a green scale). Heatmap depicting their tissue expression according to data available at Arabidopsis eFP Browser (Developmental Map dataset). Numbers inside the black boxes indicate the highest transcript level in each tissue; the gray scales are relative to these values. Striped panels indicate that no data is available.

**Figure. S4. Tissue-specific expression and light-mediated expression fold of the 41 PKs abundantly expressed in etiolated seedlings.** Left panel: PKs are grouped according to a phylogeny inferred from full-length protein alignments (AGI codes are indicated in brackets). Middle panel: heatmap depicting *PK* tissue expression according to data available at Arabidopsis eFP Browser (Developmental Map dataset). Expression levels of the top-ranked *PK* transcripts in each tissue are indicated by the numbers within black boxes; the gray scale in each tissue is relative to these values. The expression of the three metabolic *PGK* genes is not shown (striped panels) because their very high transcript levels (1000 to 3000) mask subtler expression differences among the remaining genes. Right panel: Fold change [log2(light/dark)] of *AtPKs* expression in etiolated seedlings exposed to different light treatments for the indicated times. W: white, B: blue, R: red light. Dataset authors are indicated in column headers.

**Figure S5.** Phylogenetic analysis of 135 phosphorylated PKs detected in etiolated seedlings. The inferred phylogeny was based on full-length protein alignments. Bootstrap support values >50 (from 100 replicates) are shown at the nodes. PK names in **bold:** high protein abundance; in gray: presence not confirmed; in brown: not detected in any proteome. (♦) indicate proteins detected in both phosphoproteomes. Prominent PK families are color-coded; selected RLK subfamilies are indicated.

**Figure S6.** Phosphorylation patterns in BSKs and BSLs **(A)**, and CKLs **(B)**. Gene names in **bold:** high protein abundance; in gray: uncertain presence. Domain architecture is depicted based on ScanProSite and TMHMM predictions: EKD, EATP binding site, ES/T PP. Disordered regions, based on PrDOS predictions, are <u>underlined</u> <u>in green</u>. pS: detected in both phosphoproteomes, pS: in only one dataset. Non-exclusive phosphopeptides are displayed with transparency. (*) Phosphorylations in an RxxS context.

**Figure. S7. Tissue-specific expression and light-mediated expression fold of the different members of the AtCPK family.** Left panel: CPKs are grouped according to a phylogeny inferred from full-length protein alignments (AGI codes are indicated in brackets). CPKs in **bold**: highly abundant, gray: uncertain presence; brown: absent. Middle panel: heatmap depicting *CPK* tissue expression according to data available at Arabidopsis eFP Browser (Developmental Map dataset). Expression levels of the top-ranked *CPK* transcripts in each tissue are indicated in numbers inside the black boxes; the gray scale in each tissue is relative to these values. Striped panels in *AtCPK27* and *AtCPK31* indicate that no data is available. Right panel: Fold change [log2(light/dark)] of *AtCPKs* expression in etiolated seedlings exposed to different light treatments for the indicated times. W: white, B: blue, R: red light. Dataset authors are indicated in column headers.

**Figure S8. CPK targets reported in the PhosPhAt 4.0 database and detected in the phosphoproteome of etiolated seedlings. (A)** Histogram showing the number of experimentally reported CPK targets (PhosPhAt 4.0 DB) corresponding to the CPK isoforms identified in etiolated seedlings. **(B)** Venn diagram of CPK phosphotargets detected in the phosphoproteomes from Arico et al. (2021) and Zander et al. (2020). Targets of CPK3 are shown in **bold**; targets of CPK9 are <u>underlined</u>. (*) indicate phosphoproteins containing a pS within the RxxS consensus motif. ACA8: autoinhibited Ca^2+^-ATPase isoform 8, ADH1: alcohol dehydrogenase 1, AMT1;1: ammonium transporter 1, BPL1: RNA-binding (RRM/RBD/RNP motifs) family protein, CINV1: cytosolic invertase 1, F2KP: fructose-2,6-bisphosphatase, GRF3/5: general regulatory factors 3/5, HA1/2: H[+]-ATPases 1/2, MS35Z: Ribosomal protein S24/S35, NIA2: nitrate reductase 2, PAPP2C: phytochrome-associated protein phosphatase type 2C, PIP3: plasma membrane intrinsic protein 3, PP2C33: Protein phosphatase 2C family protein, Rem1.3: Remorin family protein, RSZ22a: RNA recognition motif and CCHC-type zinc finger domains containing protein, SAPX: stromal ascorbate peroxidase.

**Figure S9. Tissue-specific expression and light-mediated expression fold of the 45 PPs expressed in etiolated seedlings.** Left panel: PPs are grouped according to a phylogeny inferred from full-length protein alignments (AGI codes are indicated in brackets). PPs in **bold**: highly abundant. Middle panel: heatmap depicting *PP* tissue expression according to data available at Arabidopsis eFP Browser (Developmental Map dataset). Expression levels of the top-ranked *PP* transcripts in each tissue are indicated by the numbers within black boxes; the gray scale in each tissue is relative to these values. Striped panels in RDO5 and PP2C80 indicate that no data is available. Right panel: Fold change [log2(light/dark)] of *AtPPs* expression in etiolated seedlings exposed to different light treatments for the indicated times. W: white, B: blue, R: red light. Dataset authors are indicated in column headers.

**Figure S10. Phosphorylation patterns in PPs.** PP names in **bold:** high protein abundance. Domain architecture is depicted based on ScanProSite and TMHMM predictions. Disordered regions, based on PrDOS predictions, are <u>underlined in green</u>. pS: detected in both phosphoproteomes, pS/pT: detected in only one dataset. (*) Phosphorylations in an RxxS context.

