## Supplementary material for "Multi-Omics Meta-Analysis Provides Insights into Reversible Phosphorylation During Arabidopsis Skotomorphogenesis": Fig. S1

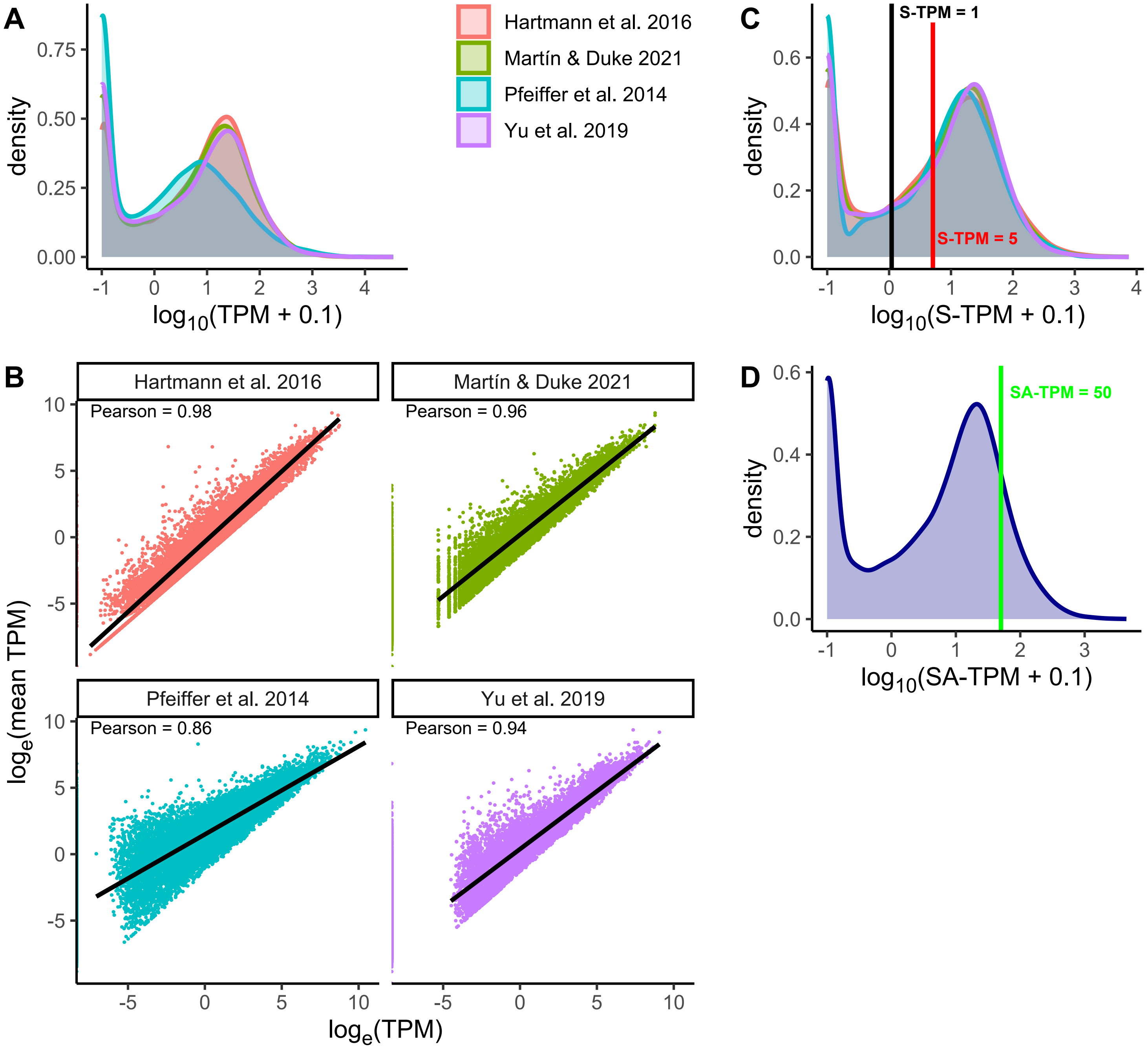


**Figure S1. Normalization of RNA-seq data from four Arabidopsis dark-grown seedlings datasets.**
**(A)** Distribution of TPM values for 27,533 genes across four datasets. **(B)** Logarithmic linear regressions for each dataset, plotting the natural logarithm of the mean TPM value per gene across datasets (y-axis) against the natural logarithm of the TPM value for the same gene in the corresponding dataset (x-axis). Pearson correlation index is indicated. **(C)** Standardized TPM values (S-TPM) were obtained by applying a regression-based transformation to each gene in each dataset. The red line indicates the threshold used to consider transcript presence (S-TPM>5 in at least three experiments). The black line indicates the threshold used to infer absence of transcription (S-TPM values < 1 in at least three experiments). **(D)** Standardized average TPM (SA-TPM) values computed as the mean of the S-TPMs across all datasets. The green line indicates the threshold for highly abundant transcripts (SA-TPM>50).
