## Supplementary material for "Multi-Omics Meta-Analysis Provides Insights into Reversible Phosphorylation During Arabidopsis Skotomorphogenesis": Fig. S2

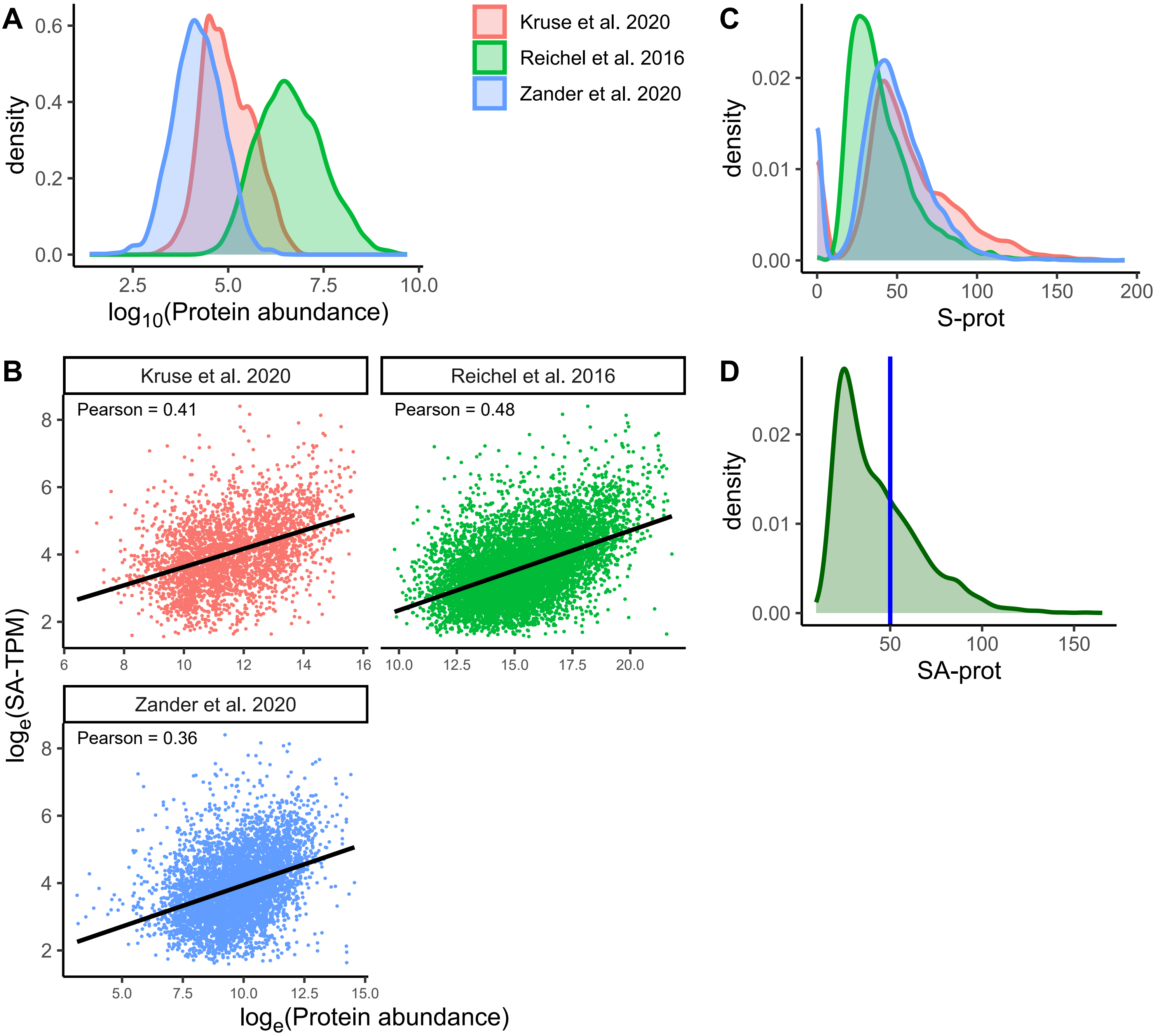


**Figure S2. Normalization of proteomic data from three Arabidopsis dark-grown seedlings datasets.** **(A)** Distribution of protein abundance across three datasets. **(B)** A linear fit on a logarithmic scale was performed between the protein abundance values in each experiment and their corresponding SA-TPM for genes with detectable transcript levels. **(C)** Distribution of the fitted values (S-prot) across the three datasets. **(D)** The fitted values were averaged to compute a standardized average protein abundance (SA-prot) for each gene. The blue line indicates the threshold for highly abundant proteins (SA-prot>50).
