## Supplementary material for "Multi-Omics Meta-Analysis Provides Insights into Reversible Phosphorylation During Arabidopsis Skotomorphogenesis": Fig. S3

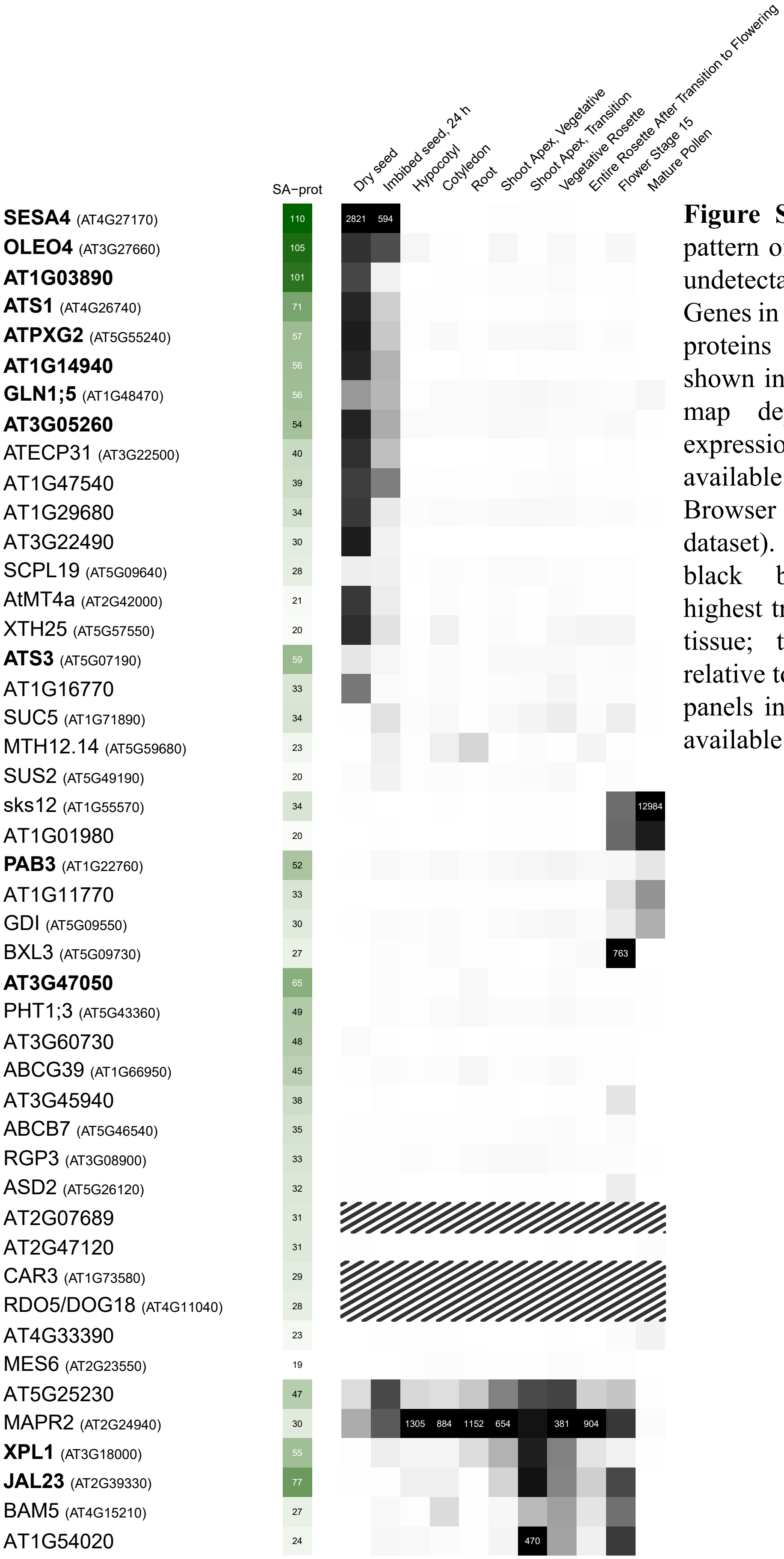


**Figure S3.** Tissue expression pattern of the 46 proteins with undetectable transcript levels. Genes in **bold**: highly abundant proteins (SA-prot values shown in a green scale). Heat-map depicting their tissue expression according to data available at Arabidopsis eFP Browser (Developmental Map dataset). Numbers inside the black boxes indicate the highest transcript level in each tissue; the gray scales are relative to these values. Striped panels indicate that no data is available.
