## Supplementary material for "Multi-Omics Meta-Analysis Provides Insights into Reversible Phosphorylation During Arabidopsis Skotomorphogenesis": Fig. S4

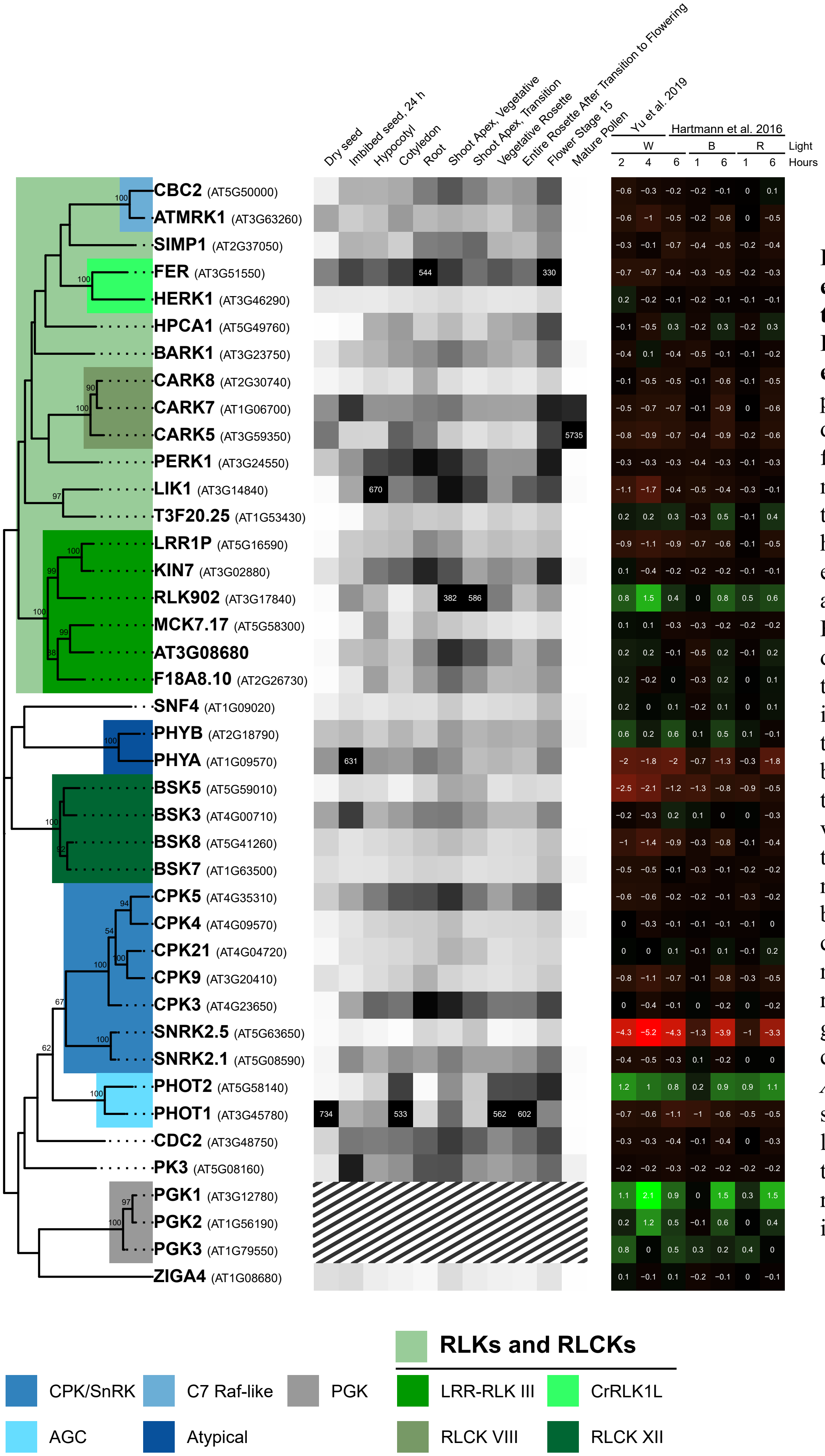


**Figure. S4. Tissue-specific expression and light-mediated expression fold of the 41 PKs abundantly expressed in etiolated seedlings.** Left panel: PKs are grouped according to a phylogeny inferred from full-length protein alignments (AGI codes are indicated in brackets). Middle panel: heatmap depicting PK tissue expression according to data available at Arabidopsis eFP Browser (Developmental Map dataset). Expression levels of the top-ranked PK transcripts in each tissue are indicated by the numbers within black boxes; the gray scale in each tissue is relative to these values. The expression of the three metabolic PGK genes is not shown (striped panels) because their very high transcript levels (1000 to 3000) mask subtler expression differences among the remaining genes. Right panel: Fold change [log2(light/dark)] of *AtPKs* expression in etiolated seedlings exposed to different light treatments for the indicated times. W: white, B: blue, R: red light. Dataset authors are indicated in column headers.
