## Supplementary material for "Multi-Omics Meta-Analysis Provides Insights into Reversible Phosphorylation During Arabidopsis Skotomorphogenesis": Fig. S5

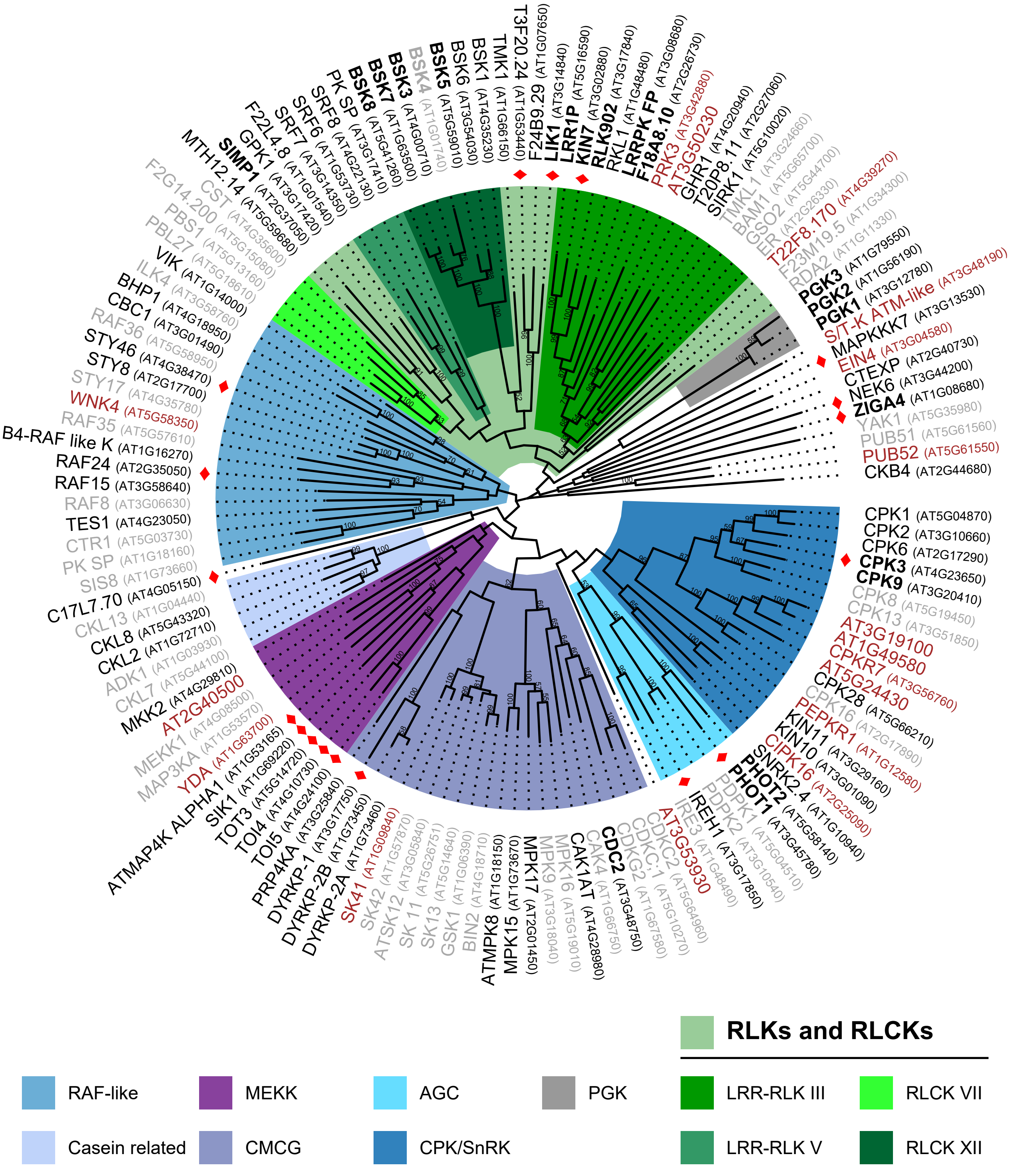


**Figure S5.** Phylogenetic analysis of 135 phosphorylated PKs detected in etiolated seedlings. The inferred phylogeny was based on full-length protein alignments. Bootstrap support values >50 (from 100 replicates) are shown at the nodes. PK names in **bold**: high protein abundance; in gray: presence not confirmed; in brown: not detected in any proteome. (♦) indicate proteins detected in both phosphoproteomes. Prominent PK families are color-coded; selected RLK subfamilies are indicated.
