## Supplementary material for "Multi-Omics Meta-Analysis Provides Insights into Reversible Phosphorylation During Arabidopsis Skotomorphogenesis": Fig. S6

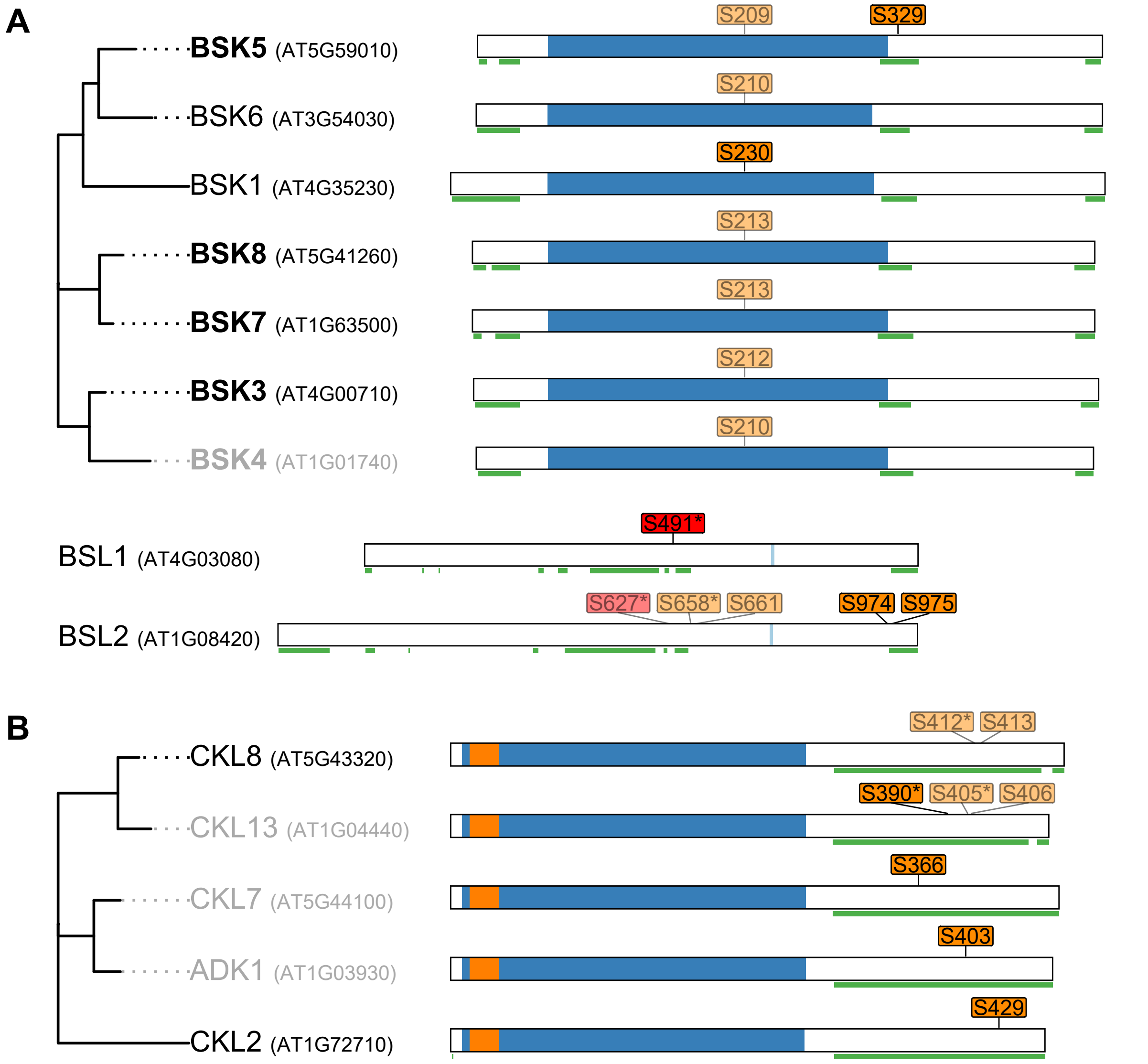


**Figure S6.** Phosphorylation patterns in BSKs and BSLs **(A)**, and CKLs **(B)**. Gene names in **bold**: high protein abundance; in gray: uncertain presence. Domain architecture is depicted based on ScanProSite and TMHMM predictions: ■KD, ■ATP binding site, ■S/T PP. Disordered regions, based on PrDOS predictions, are underlined in green. pS: detected in both phosphoproteomes, pS: in only one dataset. Non-exclusive phosphopeptides are displayed with transparency. (*) Phosphorylations in an RxxS context.
