## Supplementary material for "Multi-Omics Meta-Analysis Provides Insights into Reversible Phosphorylation During Arabidopsis Skotomorphogenesis": Fig. S7

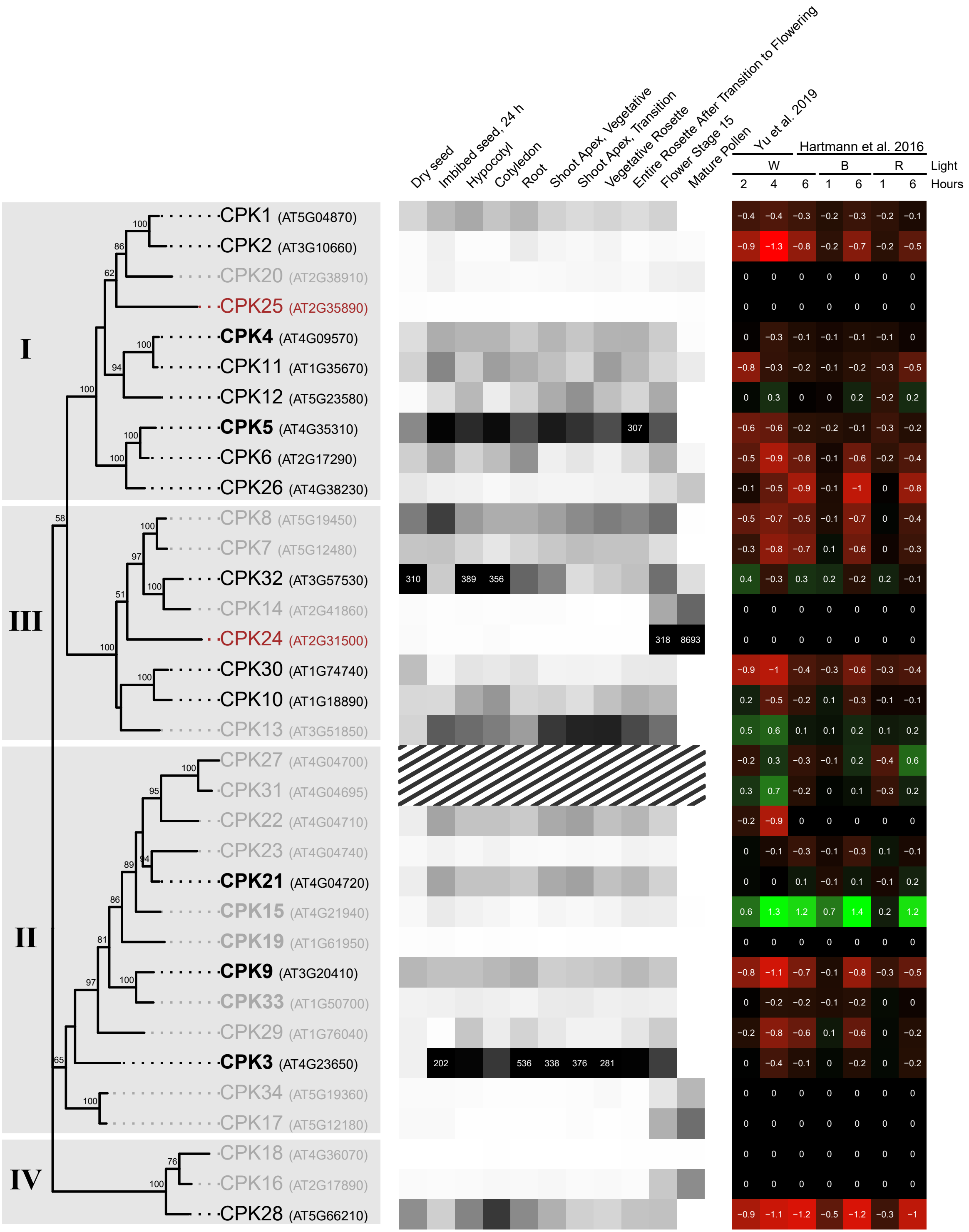


**Figure. S7. Tissue-specific expression and light-mediated expression fold of the different members of the AtCPK family.** Left panel: CPKs are grouped according to a phylogeny inferred from full-length protein alignments (AGI codes are indicated in brackets). CPKs in **bold**: highly abundant, gray: uncertain presence; brown: absent. Middle panel: heatmap depicting *CPK* tissue expression according to data available at Arabidopsis eFP Browser (Developmental Map dataset). Expression levels of the top-ranked CPK transcripts in each tissue are indicated in numbers inside the black boxes; the gray scale in each tissue is relative to these values. Striped panels in *AtCPK27* and *AtCPK31* indicate that no data is available. Right panel: Fold change [log2(light/dark)] of *AtCPKs* expression in etiolated seedlings exposed to different light treatments for the indicated times. W: white, B: blue, R: red light. Dataset authors are indicated in column headers.
