## Supplementary material for "Multi-Omics Meta-Analysis Provides Insights into Reversible Phosphorylation During Arabidopsis Skotomorphogenesis": Fig. S8

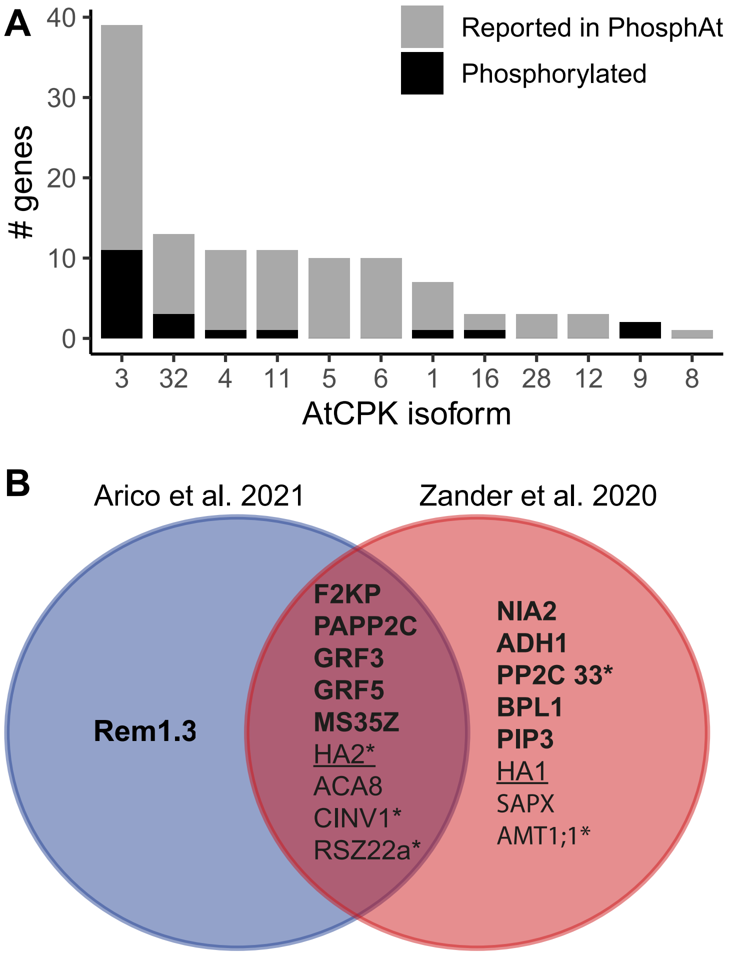


**Figure S8. CPK targets reported in the PhosPhAt 4.0 database and detected in the phosphoproteome of etiolated seedlings. (A)** Histogram showing the number of experimentally reported CPK targets (PhosPhAt 4.0 DB) corresponding to the CPK isoforms identified in etiolated seedlings. **(B)** Venn diagram of CPK phosphotargets detected in the phosphoproteomes from Arico et al. (2021) and Zander et al. (2020). Targets of CPK3 are shown in **bold**; targets of CPK9 are <u>underlined</u>. (*) indicate phosphoproteins containing a pS within the RxxS consensus motif. ACA8: autoinhibited $Ca^{2+}$-ATPase isoform 8, ADH1: alcohol dehydrogenase 1, AMT1;1: ammonium transporter 1, BPL1: RNA-binding (RRM/RBD/RNP motifs) family protein, CINV1: cytosolic invertase 1, F2KP: fructose-2,6-bisphosphatase, GRF3/5: general regulatory factors 3/5, HA1/2: H[+]-ATPases 1/2, MS35Z: Ribosomal protein S24/S35, NIA2: nitrate reductase 2, PAPP2C: phytochrome-associated protein phosphatase type 2C, PIP3: plasma membrane intrinsic protein 3, PP2C33: Protein phosphatase 2C family protein, Rem1.3: Remorin family protein, RSZ22a: RNA recognition motif and CCHC-type zinc finger domains containing protein, SAPX: stromal ascorbate peroxidase.
