## Supplementary material for "Multi-Omics Meta-Analysis Provides Insights into Reversible Phosphorylation During Arabidopsis Skotomorphogenesis": Fig. S9

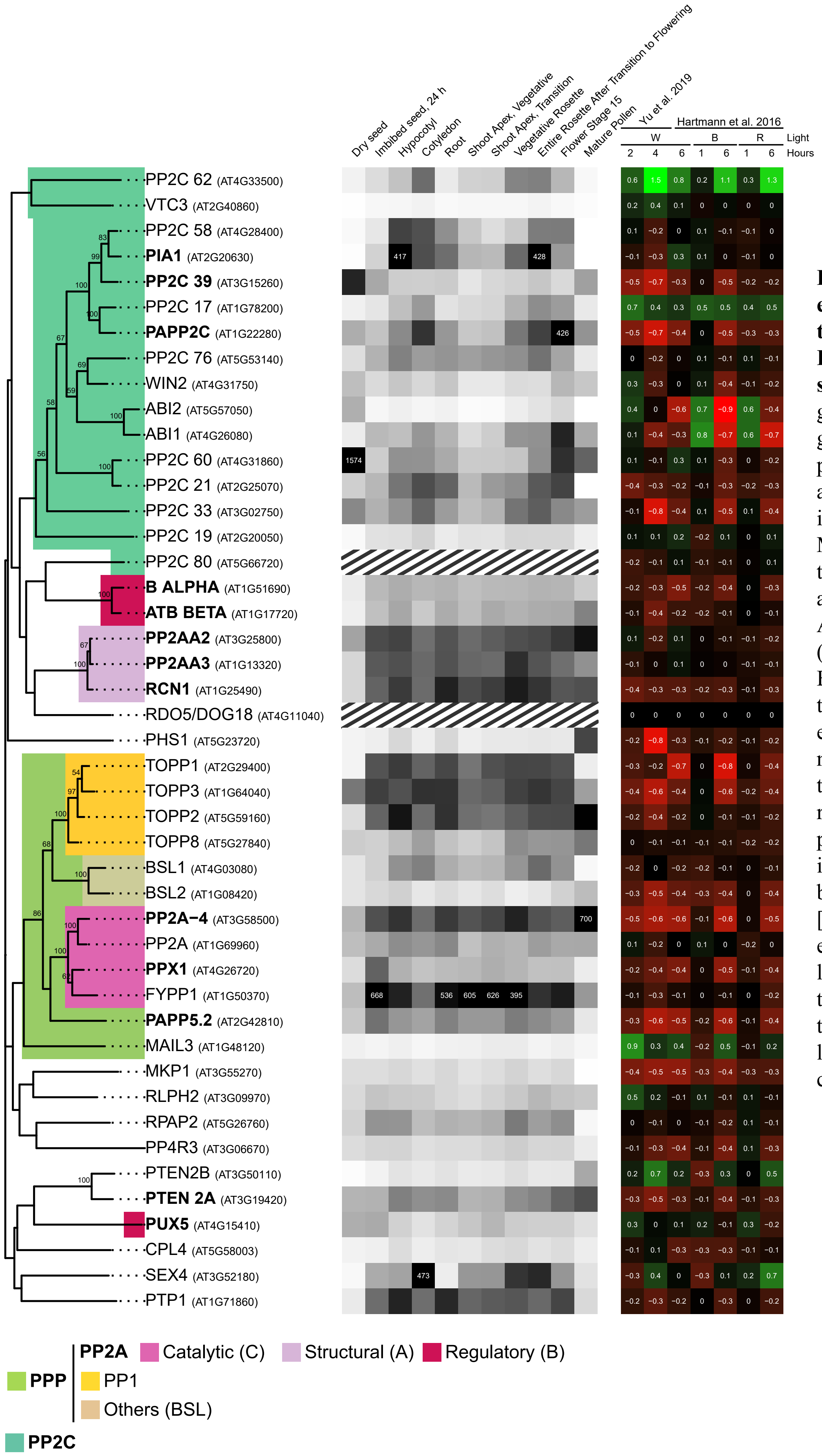


**Figure S9. Tissue-specific expression and light-mediated expression fold of the 45 PPs expressed in etiolated seedlings.** Left panel: PPs are grouped according to a phylogeny inferred from full-length protein alignments (AGI codes are indicated in brackets). PPs in **bold**: highly abundant. Middle panel: heatmap depicting PP tissue expression according to data available at Arabidopsis eFP Browser (Developmental Map dataset). Expression levels of the top-ranked PP transcripts in each tissue are indicated by the numbers within black boxes; the gray scale in each tissue is relative to these values. Striped panels in RDO5 and PP2C80 indicate that no data is available. Right panel: Fold change [log2(light/dark)] of *AtPPs* expression in etiolated seedlings exposed to different light treatments for the indicated times. W: white, B: blue, R: red light. Dataset authors are indicated in column headers.
