## Supplementary material for "Multi-Omics Meta-Analysis Provides Insights into Reversible Phosphorylation During Arabidopsis Skotomorphogenesis": Fig. S10

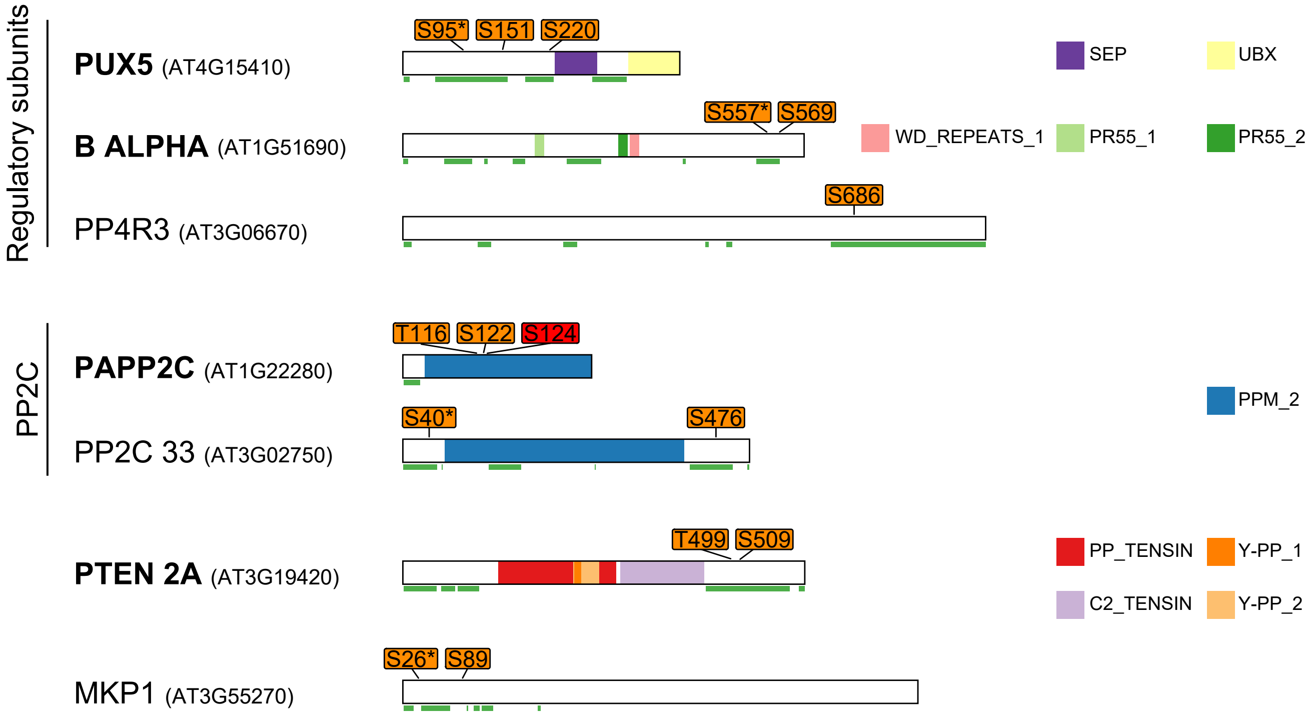


**Figure S10. Phosphorylation patterns in PPs.** PP names in **bold**: high protein abundance. Domain architecture is depicted based on ScanProSite and TMHMM predictions. Disordered regions, based on PrDOS predictions, are underlined in green. pS: detected in both phosphoproteomes, pS/pT: detected in only one dataset. (*) Phosphorylations in an RxxS context.
